# Optimizing Phylogenomics with Rapidly Evolving Long Exons: Comparison with Anchored Hybrid Enrichment and Ultraconserved Elements

**DOI:** 10.1101/672238

**Authors:** Benjamin R. Karin, Tony Gamble, Todd R. Jackman

## Abstract

Marker selection has emerged as an important component of phylogenomic study design due to rising concerns of the effects of gene tree estimation error, model misspecification, and data-type differences. Researchers must balance various trade-offs associated with locus length and evolutionary rate among other factors. The most commonly used reduced representation datasets for phylogenomics are ultraconserved elements (UCEs) and Anchored Hybrid Enrichment (AHE). Here, we introduce Rapidly Evolving Long Exon Capture (RELEC), a new set of loci that targets single exons that are both rapidly evolving (evolutionary rate faster than *RAG1*) and relatively long in length (greater than 1,500 bp), while at the same time avoiding paralogy issues across amniotes. We compare the RELEC dataset to UCEs and AHE in squamate reptiles by aligning and analyzing orthologous sequences from 17 squamate genomes, composed of ten snakes and seven lizards. The RELEC dataset (179 loci) outperforms AHE and UCEs by maximizing per-locus genetic variation while maintaining presence and orthology across a range of evolutionary scales. RELEC markers show higher phylogenetic informativeness than UCE and AHE loci, and RELEC gene trees show greater similarity to the species tree than AHE or UCE gene trees. Furthermore, with fewer loci, RELEC remains computationally tractable for full Bayesian coalescent species tree analyses. We contrast RELEC to and discuss important aspects of comparable methods, and demonstrate how RELEC may be the most effective set of loci for resolving difficult nodes and rapid radiations. We provide several resources for capturing or extracting RELEC loci from other amniote groups.

## Introduction

Though large phylogenomic datasets have become relatively easy to obtain in recent years and have led to many highly resolved phylogenetic estimates, it has become clear that the shear quantity of sequence data that can now be gathered will not unambiguously resolve some of the most difficult nodes in the tree of life. These difficulties may be caused by a number of factors including systematic error from non-phylogenetic signal or model inadequacy (Hahn and Nakhleh 2016; Reddy et al. 2017), gene tree estimation error from insufficient phylogenetic signal (Blom et al. 2017), or from natural processes such as incomplete lineage sorting and introgression (Maddison 1997; Edwards 2009) and positive selection (Castoe et al. 2009). Even as whole genomes have become easier to sequence for phylogenomics, they must still be subsetted to make aligned sets of orthologous loci. Appropriate marker selection is therefore a critical part of phylogenomics, and it is still under considerable debate what kinds of markers are the best for resolving difficult branches at various evolutionary depths. For example, questions remain whether to use coding or noncoding sequence data (Chen et al. 2017; Reddy et al. 2017), conserved or highly variable loci (Salichos and Rokas 2013; Betancur-R et al. 2014), long or short alignments (Edwards et al. 2016; Springer and Gatesy 2016), single nucleotide polymorphisms alone (SNP) or full sequence alignments (Leaché and Oaks 2017), or other types of markers that may be relatively free of homoplasy (e.g., indels, Simmons and Ochoterena 2000; SINEs, Ray 2007; transposable elements, Han et al. 2011; micro RNAs, Tarver et al. 2013). Each kind of marker possesses different trade-offs of phylogenetic information content (PIC), maintenance across evolutionary scales, susceptibility to error, and computational tractability, and these factors must be balanced during marker selection. A universal marker type for all kinds of phylogenetic questions is unlikely to exist, which may necessitate filtering for question-specific markers (Chen et al. 2015) as just a small number of loci out of thousands may have the power to resolve specific questions or drive a contentious pattern (Brown and Thomson 2017; Shen et al. 2017). Therefore, careful marker selection before sequencing to prioritize signal and resolving power over shear quantity of data may be an important step forward in phylogenomics.

### Marker Selection Trade-offs

A main trade-off in marker selection is to maximize per-locus phylogenetic signal without suffering alignment quality or excessive substitution saturation. In selecting markers, PIC can be improved either by increasing the length of the locus or by choosing loci with a faster rate of evolution. Both of these strategies increase the number of phylogenetically informative sites that can increase phylogenetic resolution (Graybeal 1994). Given sufficient time substitution saturation will eventually erode phylogenetic signal in all types of markers (Graybeal 1994), but this will occur fastest on loci that evolve more rapidly (Yang 1998). At shallow evolutionary scales it is generally agreed that rapidly evolving markers are the most effective for resolving trees because substitution saturation is unlikely to have occurred and variable markers are more likely to contain phylogenetic signal for relationships with short internodes (Dornburg et al. 2017). On the other hand, it is still under considerable debate whether conserved or variable markers are more effective at resolving deeper relationships (Philippe et al. 2011; Salichos and Rokas 2013; Betancur-R et al. 2014), as one must balance the depth of the relationships, the rate of character change affecting the chance that the signal will be masked, and the internode length since conserved loci may not have undergone any substitutions along a short branch (Townsend et al. 2012). A marker with a particular rate can have high phylogenetic utility when internodes are long, but be positively misleading when short (Dornburg et al. 2017).

Choosing longer loci will also lead to increased PIC, but one must balance another trade-off: short loci are more likely to suffer from gene tree estimation error (GTEE) due to low signal-to-noise ratio (Betancur-R et al. 2014) while long loci are more likely to carry past recombination events with different parts holding different genealogical histories (Degnan and Rosenberg 2009), and both issues are statistically inconsistent under the multi-species coalescent (Arcila et al. 2017). GTEE has emerged as a major potential pitfall in phylogenomics with summary coalescent species tree methods being particularly susceptible (Gatesy and Springer 2014; Mirarab et al. 2014a; Roch and Warnow 2015). Summary coalescent species tree methods are the most commonly used application of the multi-species coalescent to phylogenomic datasets due to computational limitations for conducting full Bayesian coalescent approaches (Mirarab et al. 2014b). The negative implications of GTEE on summary methods has prompted the implementation of tactics such as binning alignments based on information content (which involves concatenating alignments that return similar topologies in the hope of obtaining more accurate “gene tree” estimates; Mirarab et al. 2014a), or limiting datasets to high-resolution genes (Chen et al. 2015). Some methods that interpret gene trees based on quartet sub-trees may be more robust to GTEE when large amounts of data (millions of SNPs or several thousands of loci) are provided, such as ASTRAL (Mirarab and Warnow 2015) or SVDquartets (Chifman and Kubatko 2014) though they should be tested more rigorously (Roch and Warnow 2015). It may be preferable to initially select markers that hold high PIC in order to directly reduce per-locus GTEE instead of post-hoc methods to reduce it. As computational resources continue to improve future studies may be able to use preferable full Bayesian coalescent species tree methods if they instead target a reduced number of particularly informative loci (Ogilvie et al. 2016). For example, StarBEAST2 (Ogilvie et al. 2017) is faster than previous methods such as *BEAST (Ogilvie et al. 2016; Posada 2016) and these methods have both been used either by subsetting loci (Blom et al. 2017) or tips (Bragg et al. 2018). These factors indicate the necessity for a shift in the current paradigm in phylogenomic data acquisition and analysis. Future phylogenomic studies should focus on sequencing fewer loci that are longer and provide greater phylogenetic signal and optimal evolutionary rates to answer specific questions (Leaché et al. 2016). These kinds of markers are the least likely to suffer from GTEE and will also provide higher resolution for estimating the species tree (Salichos and Rokas 2013; Mirarab et al. 2014a; Roch and Warnow 2015).

Another important consideration in marker selection is the difference between protein-coding and noncoding sequences, as in some systems each data type has produced quite different phylogenetic estimates (Lavoué et al. 2003; Nikolaev et al. 2007; Shaw et al. 2007; Reddy et al. 2017). Natural selection on noncoding regions can range from highly purifying (e.g., ultraconserved elements, Katzman et al. 2007) to relatively neutral (e.g., introns, Prychitko and Moore 1997; Chamary et al. 2006). Of markers that can be confidently determined to be orthologous, introns may have the fastest rate of evolution and have been used with success to produce well-resolved phylogenies (Jarvis et al. 2014; Chen et al. 2017), yet alignments can be problematic for distantly related taxa as evolution is relatively unconstrained (Li et al. 2017). Coding sequences also evolve under a broad range of selective regimes, but are likely to always undergo some purifying selection on the resulting protein structure (Graur and Li 2000). For example frame-shifts from indels causing premature stop codons are likely to be highly deleterious. Furthermore, codon positions can evolve at substantially different rates due to their propensity for synonymous versus nonsynonymous mutations, though substitutions even in more neutrally evolving third codon positions may still be biased. In the avian phylogeny, initial concerns about the utility of coding data were that genomic-scale molecular convergence in coding sequences due to GC-biased gene conversion or other means may bias phylogenetic estimates (Jarvis et al. 2014). Instead, it is most likely that model inadequacy directly associated with at least one of the data types explains the different estimates (Reddy et al. 2017). Still, more complex models of nucleotide evolution that incorporate additional parameters for such observed differences as individual codon substitution rates (Yang and Nielson 2002), corresponding protein structures (Whelan and Goldman 2001), or biased conversion at synonymous sites (Galtier and Gouy 1998; Holland et al. 2013) are likely to improve model fit and phylogenetic estimates for coding sequences (Dornburg et al. 2017).

### Phylogenomic Dataset Types

Several targeted hybrid enrichment datasets have been developed for phylogenomics (Faircloth et al. 2012; Lemmon et al. 2012; Singhal et al. 2017) that enable researchers to capture the same sets of markers across all taxa of interest and exclude repetitive or otherwise phylogenetically misleading parts of the genome (such as pseudogenes and paralogs). The benefit of consistently using the same sets of markers across studies is that it will eventually allow for meta-analyses as more data accumulates (Lemmon et al. 2012; Streicher and Wiens 2017). Within amniotes, the most commonly used reduced representation datasets for phylogenomics are ultraconserved elements (UCEs, Faircloth et al. 2012) and Anchored Hybrid Enrichment (AHE, Lemmon et al. 2012), both of which were developed to target the variable flanking regions surrounding highly-conserved anchor points, and we describe these in more detail below. Conserved nonexonic elements (CNEEs; Edwards et al. 2017) are another recently-proposed reduced representation dataset, though these loci may suffer from gene trees with low per-locus bootstrap support when compared to UCEs and introns and their utility has yet to be rigorously tested.

Transcriptome data itself can be used for phylogenomics (Figuet et al. 2014; Wickett et al. 2014; Brandley et al. 2015), however, in order for datasets to be consistent the transcriptomes must be gathered from the same tissue-types and when analyzing transcriptome data it can be difficult to accurately assess orthologs and align different isoforms. Finally, exon capture is commonly used to sequence orthologous exons across taxa for phylogenomics, though the loci are usually not consistent across studies, as researchers usually build unique probes for their focal group. In non-model organisms, exon capture probes are usually designed by first sequencing a transcriptome (TBEC; Bi et al. 2012), but it can be difficult to determine intron-exon boundaries for TBEC probe design, which in some cases could reduce capture efficiency (but see Portik et al. 2016).

UCEs are regions of the genome (≥100 bp in length) that have high sequence identity (≥80%) across extremely divergent taxa (Faircloth et al. 2012). UCEs are often in non-coding genomic regions, though some UCEs correspond to exons (Bejerano et al. 2004). A unifying functional role of UCEs is not fully understood (Harmston et al. 2013), though they have been shown to often play a role in gene regulation (Lenhard et al. 2003; Woolfe et al. 2005; Ni et al. 2007; Warnefors et al. 2016) and development (Dickel et al. 2018) and are undergoing purifying selection about three times higher than coding regions (Katzman et al. 2007). The UCE tetrapod probeset includes approximately 5000 loci, captured using 120 bp probes (Faircloth et al. 2012). The AHE vertebrate dataset targets a much smaller number of loci compared to UCEs (400–500 depending on the study), similarly focusing on regions of the genome that are conserved across vertebrates (but not as high identity as UCEs) and flanked by more variable regions (see Lemmon et al. 2012 for additional criteria). Over 90% of AHE probe regions correspond to exons, however the flanking regions include a higher proportion of introns or other genomic elements (see Fig. 2 of Lemmon et al. 2012). In both of these sets, the majority of the phylogenetically informative sites are expected to exist in the flanking regions rather than the more conserved probe regions, in theory to allow for the most effective capture during hybridization. However, since its inception (Lemmon et al. 2012), AHE has shifted from this “anchor” method towards tiling probes across a substantially longer target region for a reduced number of loci (Prum et al. 2015; Ruane et al. 2015), highlighting the advances in sequence capture technology allowing for hybridization to highly diverged sequences (e.g., Li et al. 2013). When employed, both the UCE and AHE datasets have often been able to resolve previously difficult nodes (e.g., Crawford et al. 2012; Crawford et al. 2015; Prum et al. 2015; Bryson et al. 2016; Streicher and Wiens 2017), though short length and/or or slow evolutionary rate may make both methods susceptible to GTEE as we show in this study.

### Rapidly Evolving Long Exon Capture

Here we introduce a set of loci optimized for high-resolution phylogenomic inference: Rapidly Evolving Long Exon Capture (RELEC). We selected these loci to maximize PIC while maintaining presence, orthology, and alignment quality across broad evolutionary scales. RELEC loci may provide the most accurate phylogenetic resolution at deep and shallow divergences, while also remaining computationally tractable. While UCEs, and AHE in a similar but less extreme extent, were developed by applying a maximum evolutionary rate cutoff within a core probe region, our RELEC approach is unique in that we apply a minimum rate cutoff across long orthologous regions in order to maximize PIC and to produce robust gene trees.

Long and rapidly evolving genes hold abundant phylogenetic signal, thus increasing the chance of producing reliable individual gene trees and decreasing stochastic error associated with short genes (Salichos and Rokas 2013; Lanier et al. 2014; Mirarab et al. 2014a; Shen et al. 2016) while retaining many other benefits of exons that have led to their continued use in phylogenetics and phylogenomics. Benefits of exons include: 1.) nucleotide evolution can be modeled more complexly than noncoding regions based upon observed rate differences between different kinds of substitutions. 2.) Protein functions are often known, which in some cases may allow for studies of phenotypic or functional differences between organisms; 3.) Aligning protein-coding regions is straightforward, and translation-based alignment algorithms are accurate (Abascal et al. 2010). No other kind of marker can be easily aligned at up to 40% sequence divergence. 4.) Indels can be accurately aligned in exons because they occur predictably in multiples of 3 bp to avoid reading frame shifts and can also be valuable phylogenetic characters on their own (Simmons and Ochoterena 2000; Luan et al. 2013).

We compared the utility of RELEC, AHE, and UCE loci within the squamate phylogeny by extracting the markers from 17 genomes spanning ~200 Ma divergence. We also provide sequence data for the RELEC loci extracted from available mammal and bird genomes, making these loci easy to implement in phylogenomic studies in any amniote group (Supplemental; https://github.com/benrkarin/RELEC). We find that the selected RELEC loci are long and highly informative and can still be accurately aligned at deep divergences, while at the same time avoiding the difficulties of orthology detection. No other set of loci for phylogenomics encompasses sequences that are as long and as variable without running into alignment problems at deeper evolutionary scales. This therefore sets up the RELEC loci to be a powerful tool for resolving recalcitrant nodes in the tree of life.

## Results and Discussion

### Assembling the RELEC Loci

By aligning and comparing exons across mammal, bird, and squamate genomes, we found 179 exons that fit the RELEC criteria (see Materials and Methods for more details). Likely due to their rapid evolutionary rate, many of the RELEC genes are poorly annotated in non-mammalian genomes, which required us to manually extract and compare each candidate locus using translated sequence tBLASTn searches. By carefully assembling the dataset in this way, we are highly confident that they all are both present among amniotes and will not present issues of paralogy that would lead to incorrect tree estimates. Still, not all lineages have the complete set of 179 long exons, often due to missing data in the genomes or in a few cases from deletions (see Table S4). For example, *ENAM* cannot be found in the chicken and painted turtle genomes, likely because these lineages do not have teeth and do not need the enamolin protein. *RIKEN* appears to be deleted in the human genome, but is present in mouse and other mammal genomes. We found only two cases of duplications within amniotes for the final set of RELEC loci, but we chose to retain them because they are highly informative and the paralogy histories are clear and easy to trace. The exceptions are the sperm receptor protein, *PKDREJ*, which shows three tandem duplicates within squamates, and *CKAP5L*, which was duplicated from *CKAP5* (which does not have a single long exon) near the reptile ancestor, as it is present in crocodiles, turtles, squamates, tuatara, and some birds, but is not present in mammals, *Xenopus*, coelacanth, gar, and zebrafish.

After extracting the loci, we used tBLASTn searches to Ensembl to confirm presence and proper annotation in other tetrapod lineages. In the human genome, all RELEC genes were correctly annotated, but non-mammalian genomes had much less accurate annotations (see Fig. 1). The chicken, painted turtle, and saltwater crocodile genomes had 81–87% of RELC loci properly annotated, while *Anolis* had only 72%, and *Xenopus* only 65%. Annotation errors included incorrectly placed intron-exon boundaries, missing data, and no annotation at all. We found 161 loci in *Xenopus*, though 11 of these are reduced in size below 1400 bp, whereas in amniotes they are all retained above 1500 bp (with a few slight exceptions just below the cutoff; see Table S4). This indicates that a subset of the RELEC loci are retained at evolutionary scales beyond amniotes or tetrapods, and RELEC orthologs can likely be found in other groups as well.

**Fig. 1.**
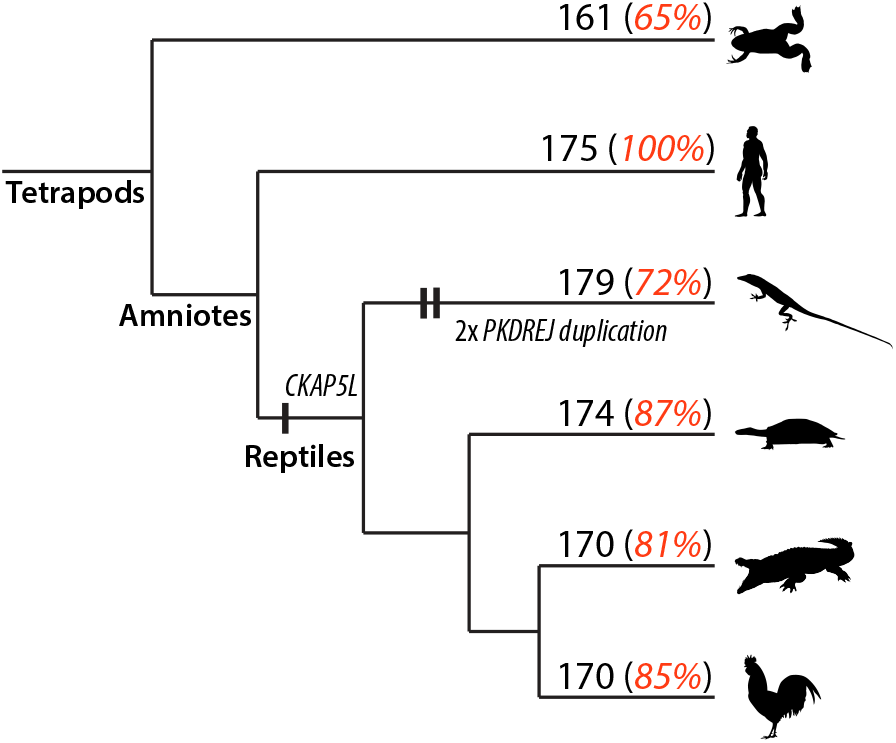
Tetrapod cladogram indicating the number of the 179 RELEC genes present in the model species for each of the main lineages, and in red the percent of them that are correctly annotated on Ensembl. Poor genome annotation outside of mammals is likely due to the rapid evolutionary rate of RELEC genes, as well as missing data in those genomes. Vertical dashes show the two exceptions where we allowed paralogs to be retained: two tandem duplications in *PKDREJ* in squamates and a duplication of *CKAP5* in reptiles. Most of the RELEC genes are present in *Xenopus*, though some are reduced in length, and many are likely present in higher vertebrate lineages as well. Silhouttes from http://phylopic.org, used under the Creative Commons (https://creativecommons.org/licenses/by/3.0/), with drawings of *Xenopus* and *Anolis* by Sarah Werning, and painted turtle by Scott Hartman.

**Fig. 2.**
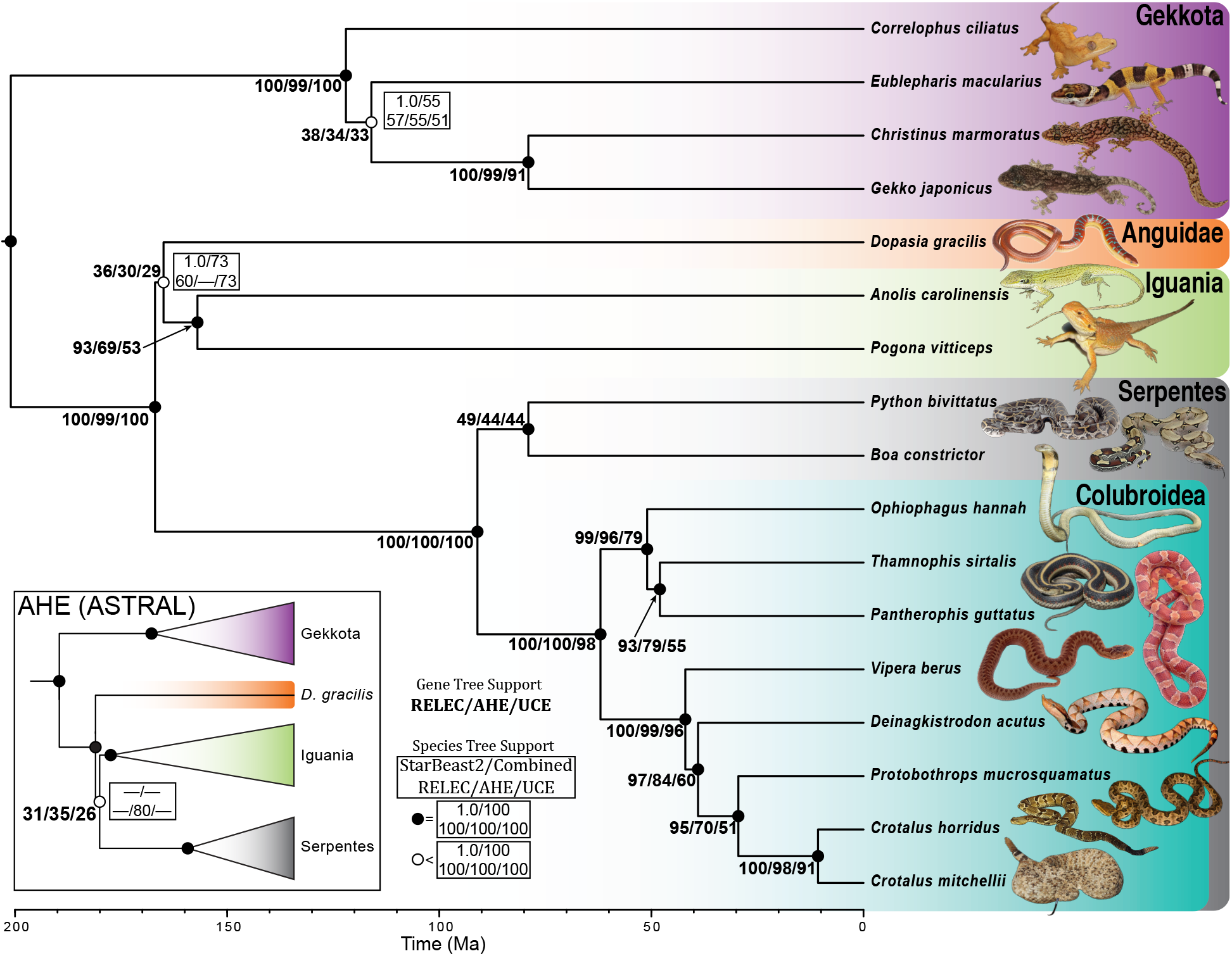
Time adjusted phylogenetic species tree for 17 squamate species using Rapidly Evolving Long Exon Capture (RELEC), Anchored Hybrid Enrichment (AHE), and Ultraconserved Element (UCE) datasets. The topology is the resulting ASTRAL species tree of the combined UCE, AHE, and RELEC dataset, which matches the UCE and RELEC ASTRAL trees as well as the StarBEAST2 species tree for RELEC. Values to the left of the node represent the proportion of maximum likelihood gene trees that support a given node (RELEC/AHE/UCE). Circles on nodes correspond to support values from ASTRAL analyses; filled circles represent 100% support for all species tree analyses (including StarBEAST2) and open circles represent any node with than 100% in any of the data sets. Values to the right of the nodes give ASTRAL support values for branches with less than 100% support. The ASTRAL topology for AHE is inset, with support for the differential node shown.

### Gene Tree-Species Tree Discordance

In this study, we estimated maximum likelihood (ML) gene trees and coalescent species trees using our newly designed RELEC dataset, and independently using the AHE and UCE datasets, as well as combined analyses of all three data-types (which we refer to as the “species tree”). All three datasets (RELEC: 179 loci, 651,434 bp; AHE: 320 loci, 427,251 bp; UCEs: 1517 loci, 1,031,286 bp; see Table 1) reconstruct the squamate phylogeny according to the species tree with minor differences at poorly supported nodes (Fig. 2; Wiens et al. 2012; Pyron et al. 2013; Zheng and Wiens 2016; Streicher and Wiens 2017). Our comparison of sequence alignments for each set show that the RELEC loci as a whole are significantly longer and contain much more parsimony informative sites than both the UCE and AHE loci (Fig. 3).

**Table 1.** General comparison of the three datasets, as acquired from the squamate genomes for this study.

| **Dataset** | **Number of Markers** | **Mean Aligned Length (SD)** | **Range Aligned Length** | **Total Number Aligned Bases** | **Completeness (% w/ 17 Taxa)** |
| --- | --- | --- | --- | --- | --- |
| RELEC | 179 | 3,639 (2,015) | 1407–12,726 | 651,434 | 98% |
| AHE | 320 | 1,335 (453) | 213–2,164 | 427,251 | 100% |
| UCEs | 1,517 | 680 (90) | 179–985 | 1,031,286 | 100% |

**Fig. 3.**
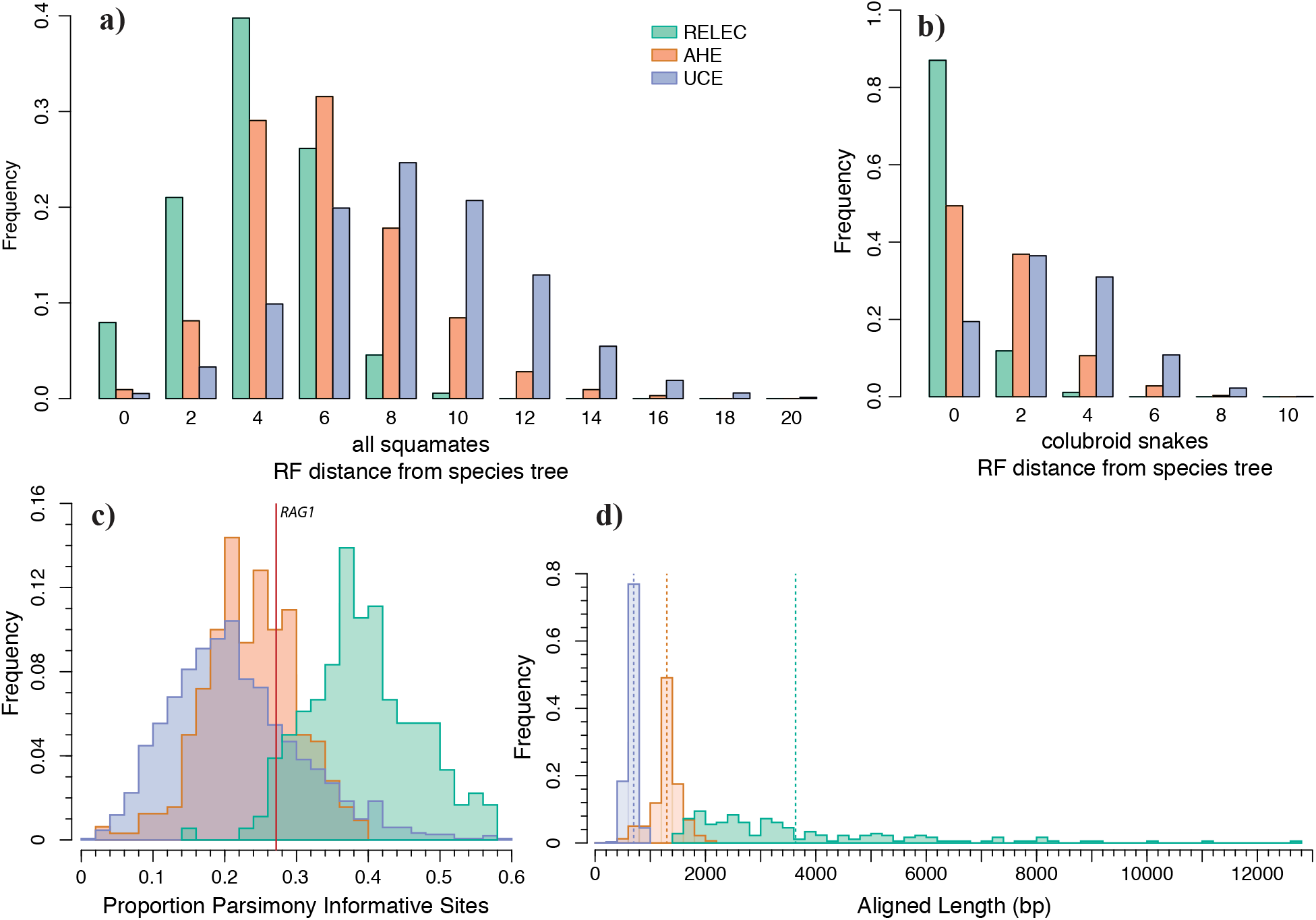
Histograms comparing features of RELEC, AHE, and UCE loci, with the y-axis in each corresponding to the proportion out of 1. (a) & (b) Robinson-Foulds distances of maximum likelihood gene trees to full squamate species tree (combined RELEC, AHE, and UCE dataset ASTRAL topology of Figure 1) and species tree limited to colubroid snakes. (c) Proportion per-locus of parsimony informative sites. Most of the RELEC genes with fewer proportion parsimony informative sites than *RAG1* (indicated by vertical red line) are genes incorporated from Wiens et al. (2012). (d) Comparison of aligned length of each locus in each dataset, with dashed vertical lines indicating the mean length in each dataset.

We first assess accuracy by comparing the ASTRAL and concatenated species tree estimates of the squamate tree for each dataset. All ASTRAL species tree nodes display strong 100% bootstrap (BS) support with two exceptions (also see open node circles in Fig. 2). 1.) Both RELEC and UCEs support *Dopasia gracilis* as sister to the iguanians, *Anolis carolinensis* and *Pogona vitticeps* (BS, RELEC: 60; UCEs: 73; Combined: 73) in agreement with the combined species tree, our RELEC StarBEAST2 estimate, and other published trees (e.g., Streicher and Wiens 2017). In contrast, both the ASTRAL and concatenated AHE trees recover support for *D. gracilis* sister to a clade composed of Iguania with snakes (BS, AHE: 80). Incomplete lineage sorting may explain the discordance on this short branch, as there are roughly equal 33% proportions for three quartet topologies at the node in each dataset (see Fig. 2), and low taxon sampling and insufficient phylogenetic signal are other possible explanations. Despite this, the RELEC StarBEAST2 analysis recovered strong (PP=1.0) support for this placement (Fig. S1), suggesting a benefit to using full Bayesian methods to co-estimate the gene trees and species tree together. 2.) Within Gekkota, the ASTRAL trees for all three datasets recover the same topology, though there exists reduced support for the placement of *Eublepharis macularius* (BS, RELEC: 57; AHE: 55; UCEs: 51; Combined: 55) and the concatenated trees showed different placements of *E. macularius* (Fig. S2). The translated RELEC amino acid data ASTRAL and concatenated trees matched the species tree exactly and showed similar support to the RELEC nucleotide analyses (Fig. S3). Separate MP-EST (Liu et al. 2010) analyses on the same sets of gene trees showed nearly identical results to those presented here (results not shown).

To assess the relative power of a gene to resolve a node at a given time period in the phylogeny, we generated phylogenetic informativeness profiles for each locus in each of the three datasets. Indeed, RELEC loci show considerably higher phylogenetic informativeness of each marker over the past 200 Ma (Fig. 4). This is due in part to their length, which is significantly correlated with informativeness (Fig. 4d), as well as their rate of evolution. RELEC loci also show a much higher proportion of parsimony informative sites than both UCEs and AHE (Fig. 3c). The informativeness profiles also show that RELEC loci are most informative at resolving younger nodes compared to AHE and UCEs (see Fig. 4e). Overall, our results show that the length and rate of RELEC loci provides them with extremely high phylogenetic informativeness across most relevant timescales and should therefore produce a greater proportion of individually reliable gene trees than do AHE and UCEs.

**Fig. 4.**
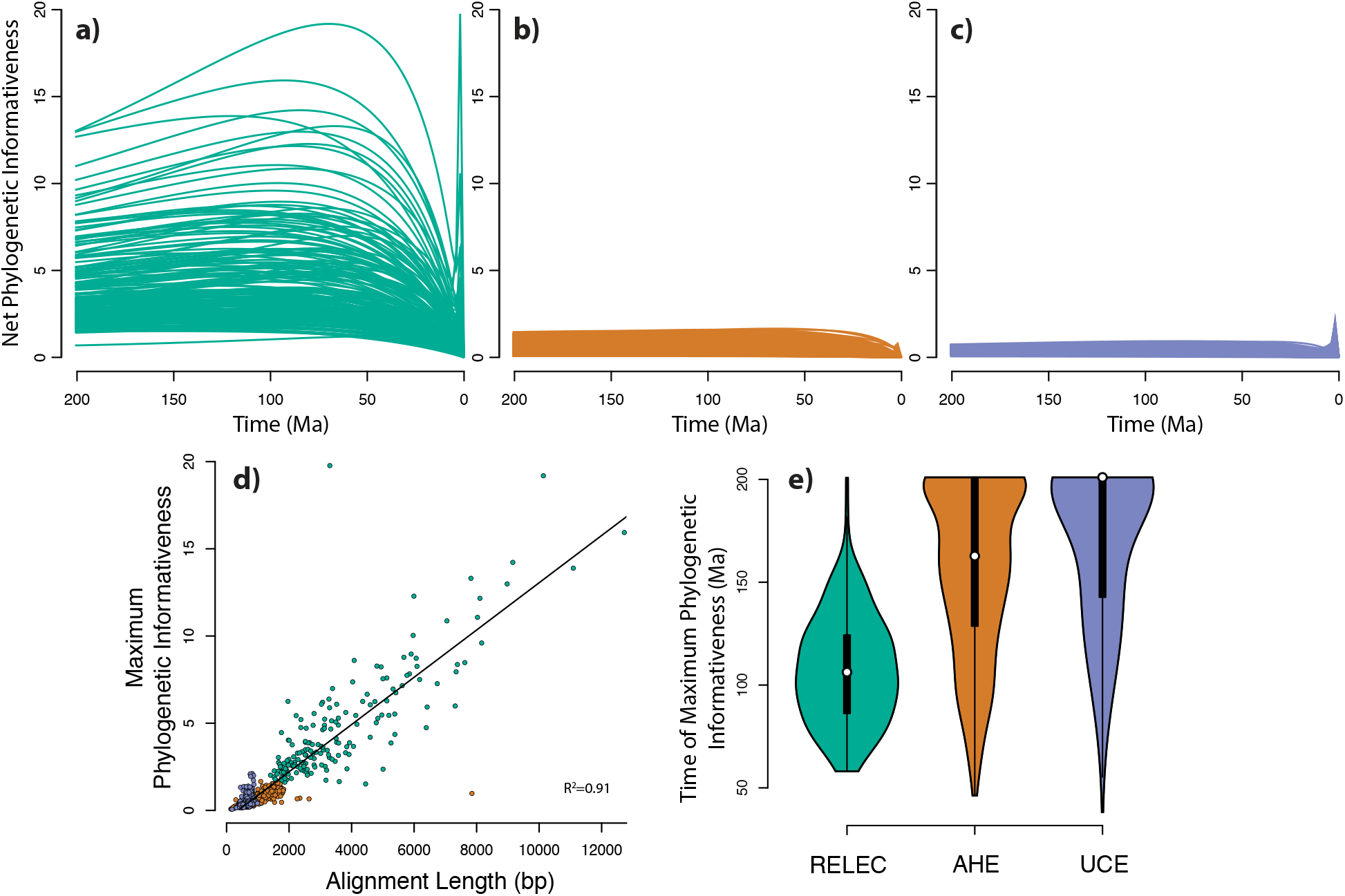
Phylogenetic informativeness of individual loci in the three datasets: (a) RELEC; (b) AHE; (c) UCEs. The y-axis is relative, and corresponds to the normalized, asymptotic likelihood that there will exist a mutation that accurately resolves a node at that point in time. The timescale corresponds to the same time-adjusted phylogeny in Figure 1. (d) The significant relationship between alignment length and the maximum value of phylogenetic informativeness reached for each locus along the curve. (e) Violin plots of the time for which each locus reaches its maximum phylogenetic informativeness, with RELEC loci optimized to have high information content at significantly younger timescales than AHE and UCEs.

We quantified GTEE between datasets using average gene tree bootstrap scores, which provide a measure of gene tree confidence (Edwards et al. 2017), and by comparing each ML gene tree to the species tree. RELEC gene trees hold significantly higher average BS support (Fig. 5b) and display higher per-node support than AHE and UCEs for all nodes in the tree that had less than 100% of the gene trees matching that node (Fig. 2). Similar to the results of Singhal et al. (2017) who compared UCEs with AHE in squamates, we also find that AHE gene trees show higher average BS support and equal or higher per-node support than UCEs (Figs. 2, 5b). RELEC and AHE performed similarly when gene trees were divided into equal-sized bins of loci analyzed with ASTRAL, with many of the jackknife replicates matching the full (dataset-specific) species tree with a bin size of ~120 loci, whereas UCEs required 500 to 1000 loci to reach a similar accuracy for the species tree estimate (Fig. 5a). The robustness of individual RELEC gene trees was especially apparent in four uncontroversial nodes in the squamate tree where 93–97% of RELEC gene trees matched the species tree for that node while AHE and/or UCEs showed substantially lower proportional accuracy (AHE: 69–84%; UCEs: 51–60%); the *Anolis-Pogona* node, the *Thamnophis-Pantherophis* node, the *Protobothrops-Crotalus* node, and the *Deinagkistrodon-Protobothrops-Crotalus* node (see Fig. 2). This suggests that RELEC gene trees are more likely to accurately resolve well-established nodes than AHE or UCEs, and this will likely apply to more difficult nodes as well. Furthermore, RELEC gene trees, on average, showed lower RF distances to the species tree than AHE or UCEs (mean RF distance, RELEC: 4, AHE, 6, UCEs: 8), indicating not only higher per-node support, but more gene trees matching the species tree in its entirety (Fig. 3a).

**Fig. 5.**
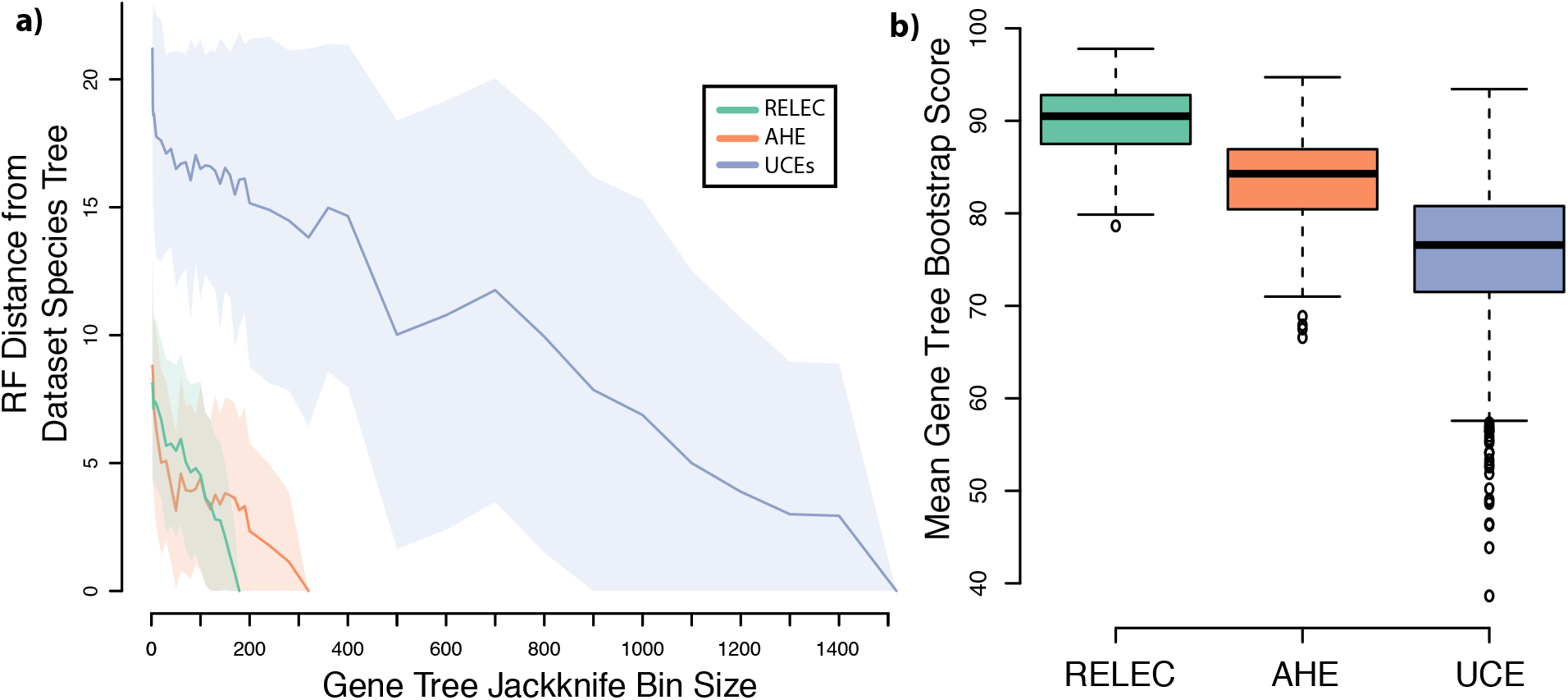
a) RF Distance of gene trees of various bin size, divided between RELEC, AHE, and UCEs. ASTRAL analyses were run on gene tree bins of increasing size with 100 random jackknife replicates each, and then compared with the species tree estimated from the largest bin. The mean RF distance is depicted by the dark line, with one standard deviation shown by the surrounding shaded area. b) Boxplots of mean bootstrap scores for ML gene trees of each dataset, with ANOVA and all comparisons significant by Tukey HSD test.

We expect that RELEC markers will be most useful at resolving more recent divergences, so we also compared gene trees to the species tree for the colubroid snakes subclade, which is the most-sampled clade in our analysis (see Fig. 2). Pyron et al. (2014) found substantial gene tree-species tree discordance among AHE loci for colubroid snakes (though these same difficult nodes are not present in this study). We find that RELEC gene trees show reduced discordance in colubroid snakes than AHE or UCE gene trees (percent of gene trees matching species tree, RELEC: 87%; AHE: 49%, UCEs: 19%; see Fig. 3b). We therefore expect that RELEC loci may more accurately resolve this clade if applied to the group as a whole, and be powerful at resolving recent radiations in general.

When examining the entire squamate tree, less than 1% of UCE or AHE gene trees match the species tree exactly, while 8.0% of RELEC gene trees match it. Though given the poor-support for the placement of *D. gracilis* in most analyses and possible incomplete lineage sorting on this short branch, if we include gene trees that differ by one node (Fig. 3b, RF=2) then a much larger proportion of gene trees are accurate (RELEC: 28.8%; AHE: 9.0%; UCEs: 3.8%). Despite this low overall rate, all three methods stand in stark contrast to the results of studies with expansive genomic-scale datasets where none of the individual gene trees match the species tree (Salichos and Rokas 2013; but see Betancur-R et al. 2014; Jarvis et al. 2014; Arcila et al. 2017). Higher discordance and GTEE in UCEs may be due to their shorter alignment lengths, lower phylogenetic signal, and less clocklike rates (Singhal et al. 2017), potentially leading to increased stochastic error for individual UCE gene trees compared to AHE or RELEC gene trees. Substitutions along short branches are less likely to have occurred for conserved loci, which may explain why relatively conserved AHE and UCE datasets have previously shown poor support for areas of the trees undergoing rapid diversification (McCormack et al. 2013; Pyron et al. 2014). As GTEE can have strong negative impacts on summary-coalescent species-tree analyses (Roch and Warnow 2015), RELEC loci that minimize it may also be the most effective for these tree-building methods.

### Utilizing Full Bayesian Coalescent Species Tree Methods

Though normally too computationally demanding to be utilized in phylogenomic-sized datasets, given the relatively small taxon-set in this study we were able to carry out a Bayesian coalescent species tree analysis on the RELEC dataset using StarBEAST2. We visualized results from both MCMC chains in Tracer 1.6 (Rambaut and Drummond 2013) and both runs reached stationarity relatively quickly, with reasonable computation times. The analysis converged on the same species tree as the combined dataset ASTRAL species tree, with posterior probabilities of 1.0 for all nodes (Fig. S1). Furthermore, there appeared to be little species-tree discordance even at poorly supported nodes in ASTRAL analyses (Fig. 2), indicating strong support for the sister relationship between Anguidae and Iguania. Given current computational power available to most researchers, we are unsure if it is possible to use StarBEAST2 for analyses of datasets much larger than this study as number of tips increases computation time rapidly (Ogilvie et al. 2016, 2017), however our results show promise for the future possibility of using these comprehensive Bayesian coalescent species tree analyses for phylogenomic datasets with fewer but longer and highly informative loci.

### Substitution Saturation

Substitution saturation decreases phylogenetic signal in sequence data by masking relevant mutations for phylogenetic inference, and has negatively affected the ability to reconstruct deep nodes in the tree of life. It is especially common in mitochondrial genes, as they evolve quickly and have a lower effective population size (Jackman et al. 1999). Researchers may be concerned that some RELEC markers may experience saturation at even moderate divergences because they evolve so rapidly, as substitution saturation has often been implicated as a problem for phylogenomics (Jeffroy et al. 2006; Parks et al. 2012; Breinholt and Kawahara 2013; Dornburg et al. 2014). However, saturation of nuclear genes normally only affects studies attempting to reconstruct deep relationships, such as the placement of turtles with respect to other reptiles (Chiari et al. 2012) and crown vertebrate lineages (Dornburg et al. 2014). RELEC loci should primarily be used to recover younger relationships within (not between) the major amniote groups, such as relationships among or within snake, gecko, or iguanian families. Within these timescales, saturation plots for each RELEC locus up to over 200 Ma divergence times overall show little evidence of substitution saturation at moderate timescales (Figs. 6, S4). In just a few cases the 3^rd^ codon position appears to show a signature of saturation by 160 Ma (e.g., *RAI1*, *PTGER4*, *FBXO34*; Figs. 6, S4), though the other codon positions appear unsaturated and are still accumulating raw pairwise distance. Even the 3^rd^ codon position of some of the most rapidly evolving genes, (e.g., *ASXL1*, *BRCA1*, *SETX*) show linearly increasing divergence up to the oldest divergence times at about 200 Ma. In comparison, the 3^rd^ codon position for mitochondrial ND2 sequences from each taxon shows rapid substitution saturation, as expected, by about 60 Ma (see Fig. 6). Given the limited evidence for saturation even at divergences spanning all squamates, we expect that most researchers will not encounter any substitution saturation in RELEC loci at the moderate evolutionary scales usually encountered in the majority of phylogenomic studies. The phylogenetic informativeness profiles suggest that RELEC loci peak in informativeness between approximately 25 and 150 Ma (Fig. 4), and we therefore recommend these scales as the regions where RELEC loci would be most effective. Still, we do not expect slight 3^rd^ codon position saturation to strongly skew phylogenetic analyses if there is sufficient signal at the other sites.

**Fig. 6.**
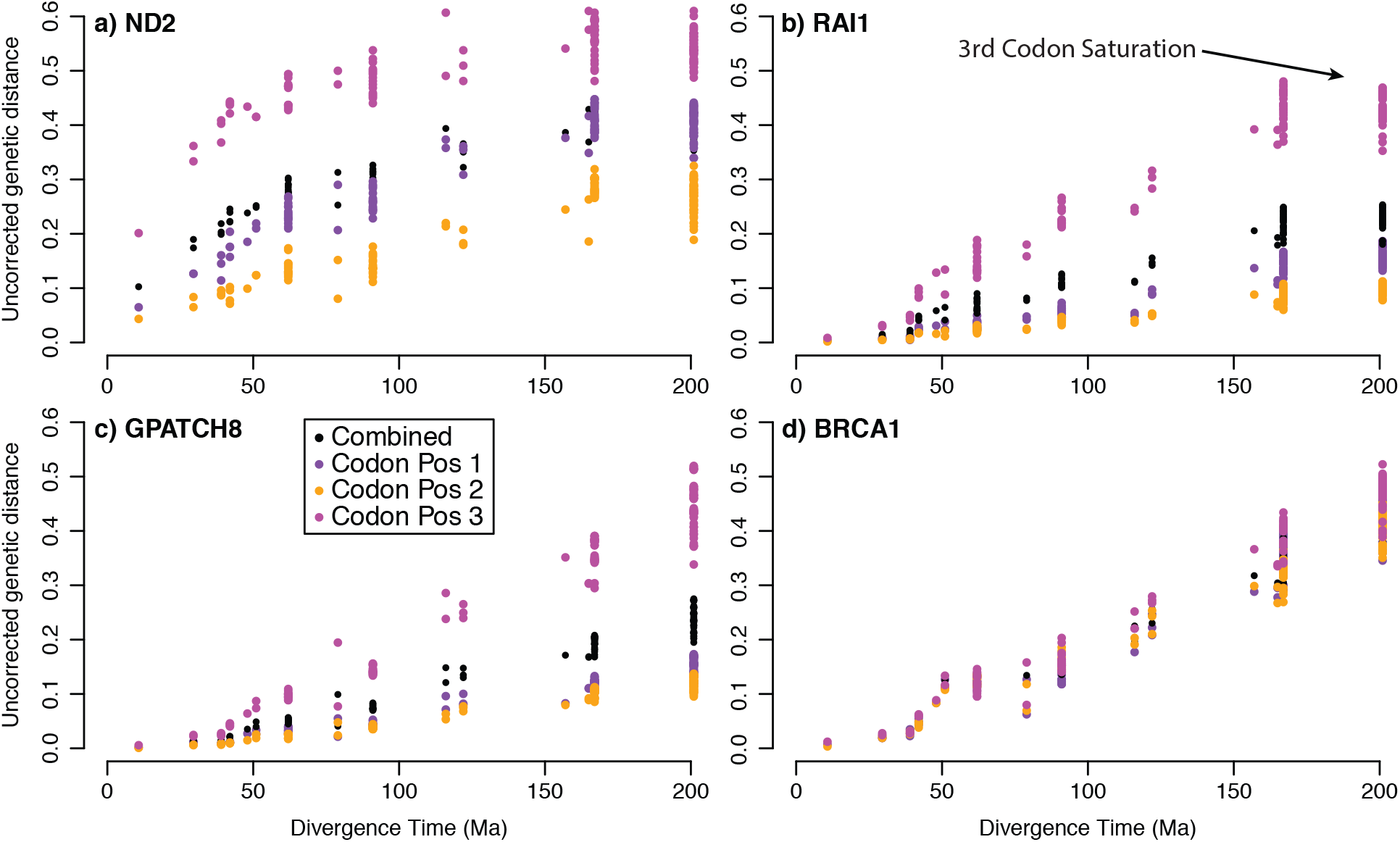
Plots to assess substitution saturation for three RELEC loci: (b) *RAI1*, (c) *GPATCH8*, (d) *BRCA1* based on comparison of raw pairwise sequence distance versus divergence time. (a) Mitchondrial *ND2* is shown for reference. A signature of saturation is present where pairwise distance does not increase despite increasing divergence time, which is prevalent in all three codon positions in *ND2*. Most RELEC loci show a pattern similar to *GPATCH8* or *BRCA1*, with a linear increase in pairwise distance over time, while *RAI1* shows some evidence of 3^rd^ codon saturation after 160 Ma (see arrow). For all saturation plots, see Supplemental Figure S4.

Exons that have roughly equal rates of evolution at all three codons are unusual (Fig 6d) but we observed this pattern for 22 RELEC exons (10%). Typically, 3^rd^ codon positions evolve much faster and this pattern is often observed across large multi-locus data sets (e.g., Baker et al. 2014 for 594 protein-coding loci). These loci may be undergoing relaxation of purifying selection and are a reflection of the increased phylogenetic utility of RELEC loci. However, for the entire RELEC dataset, the difference in slopes between 3^rd^ position and 1^st^ and 2^nd^ positions is not significantly correlated with the RF distance from the species tree as might be expected if equal rates at all positions were associated with more accurate gene trees.

### Recombination and Linkage

The chance of a gene carrying a past recombination event increases for longer genes and if recombination occurs can lead to neighboring segments of DNA with different genealogical histories that can skew phylogenetic estimates (Posada and Crandall 2002; Degnan and Rosenberg 2009). A benefit of using UCEs or RADseq is that one may gather large numbers of loci that can be analyzed independently with one SNP per locus, disregarding potential linkage disequilibrium (Leaché and Oaks 2017), but this method would not be appropriate if only capturing a small number of RELEC loci. Transcriptome data with loci that span multiple distant exons are the most likely kind of data to be susceptible to intragenic recombination (Springer and Gatesy 2016) that do indeed violate assumptions of the multi-species coalescent (Edwards et al. 2016). RELEC loci span from approximately 1,500 to 12,000 bp in aligned length (Table 1), so it is possible that recombination could occur within them, albeit with lower frequency than with transcriptome data that can span hundreds of thousands of bases. Still, we do not expect recombination to be a severe problem for RELEC as recombination events leading to incomplete lineage sorting appear to be rare in real-world datasets (Edwards et al. 2016), and unrecognized intralocus recombination events have been shown to have little effect on summary-coalescent species tree analyses (Lanier and Knowles 2012). We hypothesize that except in rare cases it is more valuable to capture longer genes that contain greater phylogenetic signal (Edwards et al. 2016) and are more likely to recover the true species tree (Fig S5; Mirarab et al. 2014a; Roch and Warnow 2015).

We assessed potential linkage of markers by comparing map distances of loci on the *Anolis* macrochromosomes (Fig. 7). For RELEC loci, only the few cases where we used two exons from a single gene were loci within 50,000 bp of each other. In more complex datasets, linkage should perhaps be assessed for these closely spaced loci, but in general we don’t expect linkage to pose a problem for RELEC. In contrast, approximately 20% of UCE loci are within 10,000 bp of each other, and more than half are within 50,000 bp. This is likely due to the sheer number of UCE probes, but this potential linkage could pose a bias if not accounted for as certain gene tree genealogies from loci in areas of tight linkage may be more common than others and affect species tree analyses. AHE loci show similar map distances as the RELEC loci, but in examining this we encountered some previously unidentified aspects of the AHE loci. The updated AHE vertebrate probes from version 1 vertebrate (Lemmon et al. 2012) to version 2 removed originally separate markers with overlapping flanking sequencing that would lead to duplicated data and also attempted to combine multiple AHE loci on a single gene into one locus (Prum et al. 2015; Ruane et al. 2015; Tucker et al. 2016), however there remain several cases of multiple AHE loci on a single gene (Table S2). For example, in version 2 we found 11 independent AHE loci representing multiple regions of a single gene, *TTN*, spanning over 240,000 bp in length. Furthermore, *FAT4* and *RHOV* each have three AHE markers representing these genes, and 12 other genes have two AHE markers associated with them. In total, 41 of the AHE version 2 vertebrate loci show duplicates, and many of the targets are within 10,000 bp of each other on the *Anolis* chromosomes (see Table S2). It is therefore important that studies utilizing AHE do not treat closely separated loci independently and are careful to account for potential linkage or functional similarity. We also found using blast searches to the www.ensembl.org database that the proportion of target regions containing coding sequence has increased from 90% in version 1 (Lemmon et al. 2012) to 98% in version 2 (this study; probes from Ruane et al. 2015), suggesting that it is now primarily a protein coding dataset yet studies have not utilized known rate differences among codon positions in evolutionary models or partitioning schemes.

**Fig. 7.**
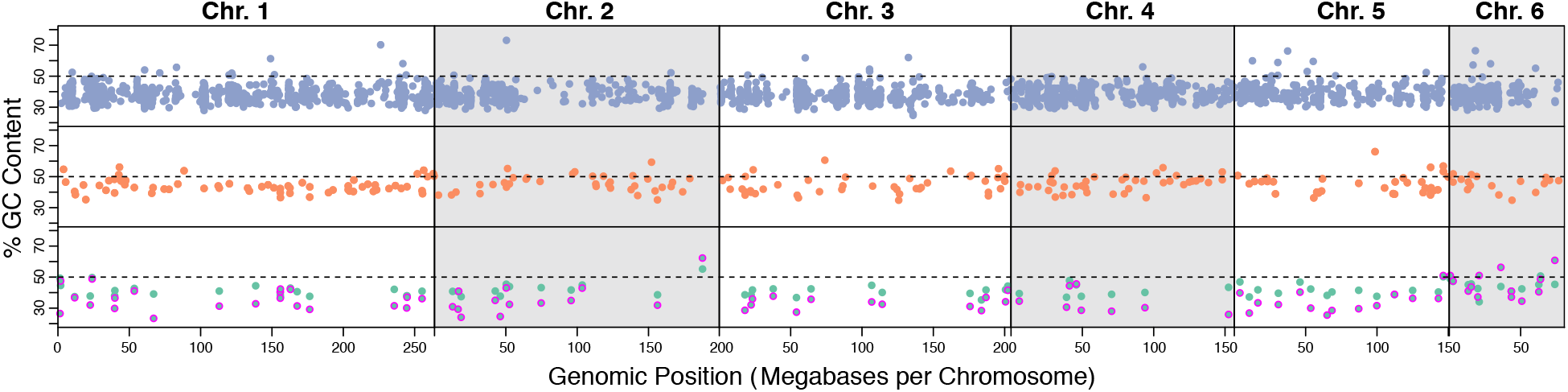
The genomic positions of UCE (top), AHE (middle), and RELEC (bottom) loci on the six *Anolis carolinensis* macrochromosomes. Markers that correspond to microchromosomes or unmapped regions are not shown. The y-axis shows the % GC content at each locus in *Anolis carolinensis*, with dots outlined in magenta in the RELEC panel corresponding to 3^rd^ codon GC content. A pattern of GC content increase at chromosome ends (where recombination is highest) would be evidence for GC-biased gene conversion, though this pattern is not apparent in any dataset.

### Coding vs. Noncoding Markers

There has been speculation that phylogenomic analysis of exons may be misleading due to biased gene conversion causing genomic-scale molecular convergence of protein-coding sequences (Jarvis et al. 2014). Jarvis et al. (2014) proposed that potentially misleading phylogenetic results in birds may be due to GC-biased gene conversion that acts most strongly in highly recombining regions, as they found rapidly evolving genes near the ends of chromosomes had increased GC content. However, Reddy et al. (2017) rejected this hypothesis by showing that the discordance between coding and non-coding datasets was likely caused by data-type effects due to violation of models. Still, the problem of GC-biased gene conversion is less likely to occur in squamates than in mammals or birds because squamates show decreased GC-content heterogeneity across chromosomes (Fujita et al. 2011) and between different lineages (Figuet et al. 2014). We plotted GC and GC3 content variation for loci between the three datasets on the *Anolis carolinensis* macrochromosomes (Fig. 7). While there does exist some variation in GC content, it is scattered across the chromosomes, and we do not observe a pattern of increased GC content (or 3^rd^ codon GC content) at the chromosome ends where recombination rates are highest and GC-biased gene conversion tends to occur. As biased gene conversion has been implicated in many systems and can negatively impact phylogenetic reconstruction (Marais 2003; Gruber et al. 2007), future studies should implement alternate models that allow for base frequency nonstationarity and should assess model fit in both coding and noncoding regions (Lockhart et al. 1994; Gowri-Shankar and Rattray 2007).

Despite concerns that selection on protein-coding regions could lead to misleading phylogenetic relationships, we emphasize that selection is not necessarily more intense on coding sequences relative to other genomic regions. For example, introns likely undergo relaxed selection (Chamary et al. 2006) while UCEs must be undergoing strong purifying selection to remain conserved across such distant groups (Katzman et al. 2007). In contrast, a distinct advantage of using protein-coding DNA sequences for phylogenetics is that nucleotide evolution can be modeled more precisely and the strength of selection can be quantified by comparing rates of synonymous and nonsynonymous mutations (McDonald and Kreitman 1991). Rate differences among 1^st^, 2^nd^, and 3^rd^ codon positions, for example, are a well-understood bi-product of selection acting more strongly against non-synonymous versus synonymous mutations (Jackman et al. 1999) and this among-site rate variation in coding sequences may be beneficial to phylogenetic reconstruction (Klopfstein et al. 2017). Translations to amino acid sequences are also commonly used for phylogenetic analysis (Fig. S3) and detailed codon models can accurately parameterize for rate differences between nonsynonymous and synonymous substitutions or between all codons (Yang 2007), and can now be rapidly implemented in recent versions of IQtree (Nguyen et al. 2015). Finally, relaxed-clock models can allow for accurate phylogenetic reconstruction by accommodating for lineage-specific evolutionary rate differences. Analyses of RELEC datasets will benefit by making use of our increased understanding of protein evolution and may allow for greater confidence in phylogenetic hypotheses.

### Biological Processes

There is unlikely to be a unifying functional role of RELEC loci, nor one single process that maintains long exons that also rapidly mutate. We searched for functional patterns among RELEC genes by investigating expression levels across human tissues (data of Fagerberg et al. 2014). We found that RELEC genes are expressed in humans at significantly different levels compared to the background of all genes (see Fig. 8; X^2^ = 27.3; p=0.00029). Specifically, the largest proportional category of RELEC genes (46.2%) are those that are expressed in low levels across all tissues, over a background level of (32.4%), while very few RELEC genes are expressed at high levels across all tissues (7.0%), compared to a higher background rate (13.8%). Furthermore, almost none (0.6%) of the RELEC genes are highly tissue enriched. These patterns suggest that many RELEC genes function as important proteins that are used in all tissue types, with low expression rates potentially loosening constraints on amino-acid substitution rates, but broad expression across all tissue types constraining the size of the longest exon and the corresponding protein. An important caveat of this analysis is that some RELEC gene isoforms (in humans) are much shorter and do not contain the long RELEC exon we selected for phylogenomics. Therefore, the isoform containing the RELEC exon may not be expressed as broadly as shorter isoforms of the gene, and the long isoform may serve a more specific function. Exceptions to these patterns are genes that are only expressed briefly in development and/or in limited tissue types. For example, *ENAM* (enamelin), encodes the largest protein of the enamel matrix and is only expressed during tooth growth (https://ghr.nlm.nih.gov/gene/ENAM). *RAG1* is involved in immune response and is only expressed in lymphocytes, and likely needs to evolve rapidly to deal help recognize different substances (https://ghr.nlm.nih.gov/gene/RAG1).

**Fig. 8.**
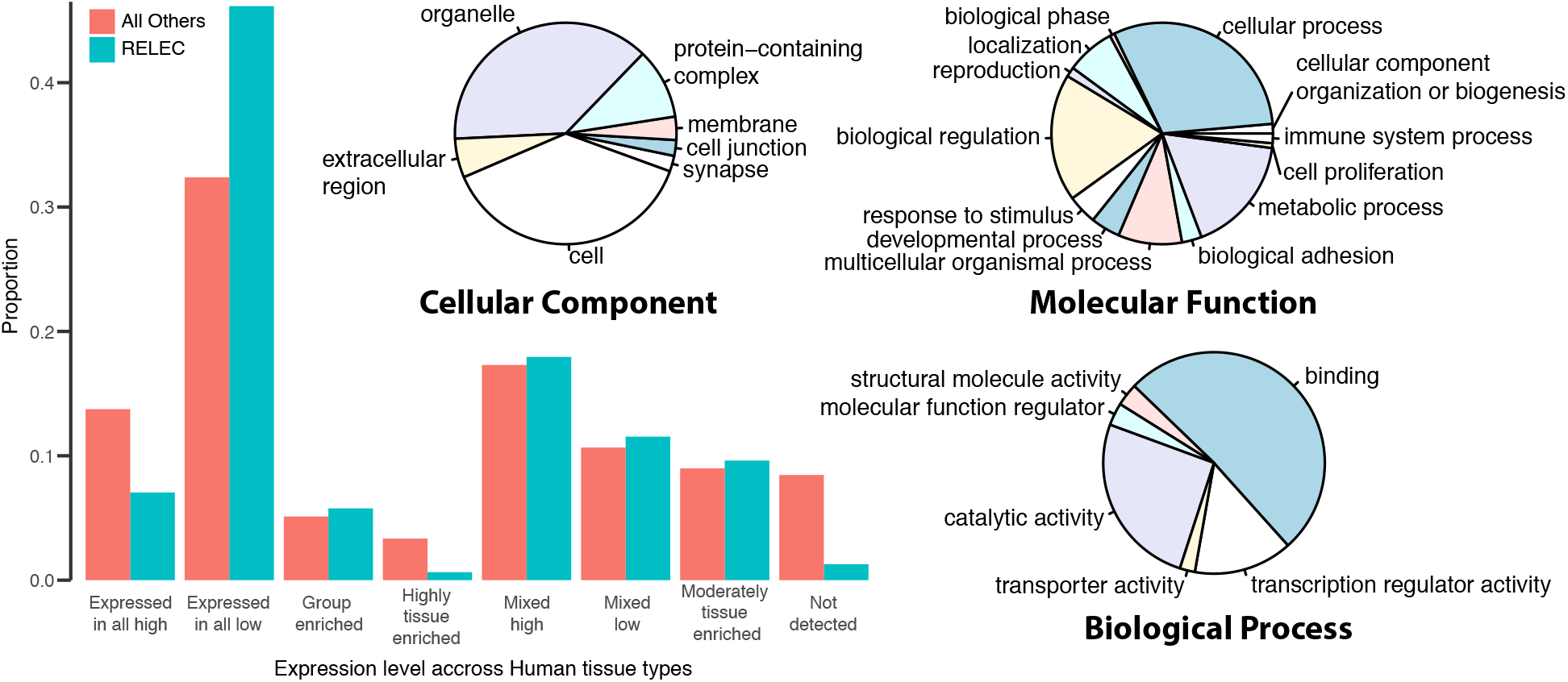
(Left) Comparison of expression level categories of RELEC genes to background levels in human tissues. RELEC genes tend to be expressed in all tissues at low rates, while fewer are expressed in all tissues at high rates (X^2^=27.3; p=0.00029). (Right) Pie charts of gene ontology (GO) functional categories for RELEC genes, divided into biological process (140 hits), molecular function (87 hits), and cellular component (90 hits), with significant enrichment for each.

We also assessed enrichment of GO functional categories of the RELEC genes as a whole on the PANTHER classification system (Fig. 8). We found significant enrichment of RELEC genes for Biological processes (140 hits), specifically for binding (51.1%) and catalytic activity (25.6%). The molecular function category (87 hits) was enriched for cellular processes (30.7%), metabolic processes (17.1%), and biological regulation (18.6%). Finally, the cellular components categories (90 hits) with the highest proportions of RELEC genes are cells (37.9%) and organelles (37.9%). The enrichment for binding suggests that the long RELEC proteins might require rapid change in amino acid sequence in order to complete their designated binding task. For example, the *BRCA1* and *BRCA2* genes are well-studied and known to be used to repair damaged DNA (Moynahan et al. 1999; Moynahan et al. 2001), and we speculate that this function may also require rapid amino acid replacement to account for different kinds of DNA damage. It is important that sets of phylogenomic loci do not all contribute to the same molecular function or pathway, as this might lead to linked evolutionary histories among loci that could bias phylogenetic estimates. The PANTHER analysis clearly shows that RELEC encoded proteins provide a wide variety of biological functions, and therefore are unlikely to be susceptible to this bias.

### Practical Implementation

Applying RELEC to any amniote group is relatively straightforward. Probes can be designed from transcriptomes or genomes and can be sequenced using many available methods (e.g., Illimuna, Fluidigm, etc.). Target enrichment for long exons among divergent taxa has been well-established in numerous studies (Bi et al. 2012; Li et al. 2013). We are currently in the process of using Illumina sequencing for RELEC loci in diplodactylid geckos using probes designed from *Correlophus ciliatus*. We also extracted the target sequences from the *Gallus gallus* genome for use in avian phylogenomics. For the geckos, we developed custom biotinylated RNA bait libraries using the MYBaits target enrichment system (MYcroarray, Ann Arbor, Michigan). Based on the 179 RELEC exons, we designed about 10,000 probes consisting of 120 bp baits with 60 bp overlap between baits, targeting exons 120 bp or larger. The baits were filtered for repetitive sequences by MYcroarray using RepeatMasker (http://www.repeatmasker.org/) and compared to the entire *Correlophus* genome to ensure unique binding of baits. Our procedure for capturing RELEC baits follows the detailed protocol for exon capture in sharks (Li et al. 2013) which captures divergent sequences by both decreasing the temperature of hybridization successively and increasing the time of hybridization of the baits to the target. The use of RELEC loci in future studies would involve designing a set of RELEC bait libraries based on the genome of a member or close relative of the group of interest. We extracted the RELEC loci from several mammal and bird genomes, and provide these sequences for users to build probes or to use them to extract RELEC loci from other genomes using custom scripts (https://github.com/benrkarin/RELEC).

It is possible that substantial genetic divergences arising from rapidly evolving markers could lead to poor hybridization efficiency in distant taxa. Though drop-off in exon capture efficiency for divergent taxa has been shown to occur in some experiments (Bi et al. 2012; Bragg et al. 2016), it is common for sequences to still be effectively captured at high enough levels for phylogenetic analysis even across extremely divergent taxa (Li et al. 2013; Ilves and López-Fernández 2014; Portik et al. 2016; Bragg et al. 2018). By coupling tiling across these relatively long probe regions and sequencing longer read lengths (e.g., MiSeq paired-end 300 bp reads), we expect to maintain capture efficiency for the RELEC markers even at substantial genetic divergences.

As it is becoming easier and less expensive to sequence full genomes or very large sets of loci, we expect that RELEC loci will most commonly be applied to new systems in two ways: 1.) targeted directly using bait capture, possibly in conjunction with other sets of loci, or 2.) extracted from whole genomes as a unique marker set for comparison with other sets. With the bait capture approach, we expect that RELEC loci will often be targeted in combination with other exons of known functional interest, and often in combination with other sets such as the Squamate Conserved Loci (SqCL) probes of Singhal et al. (2017), which combine AHE, UCEs and other commonly used squamate markers (Wiens et al. 2012), and CNEEs (Edwards et al. 2017). If a strongly supported phylogeny is the primary goal of a study, these reduced representation datasets are the most cost-effective strategy. If the researcher has other questions, such as phenotype-genotype associations, genome architecture, etc., then whole genomes can provide the means to investigate these topics and also extract different loci sets, such as RELEC, for estimating the phylogeny of the group. When whole genomes are sequenced, RELEC loci may be used as a preferred set that can be run as a whole using intensive full Bayesian coalescent species tree methods that are powerful at dealing with incomplete lineage sorting, as we have done in this study.

### Conclusion

The recent acceleration in the generation of full genomes has allowed for the development of unique sets of markers that can be tailored for particular research questions in non-model organisms. RELEC is one such example of a set of loci primarily intended to be capable of resolving difficult nodes in the tree of life, focusing on maximizing phylogenetic signal while maintaining orthology across evolutionary scales and remaining computationally tractable for comprehensive phylogenetic analyses. In the future, choosing an appropriate strategy for selecting sets of sequences to make reliable gene trees will become increasingly important. As more whole genomes are sequenced and published, our approach to finding and utilizing long stretches of informative and comparable coding sequences can be used effectively to generate a set of reliable gene trees. We look forward to future phylogenomic studies that will assess the utility of RELEC loci at resolving difficult nodes.

## Materials and Methods

We generated and tested the set of RELEC markers for an amniote dataset under the following criteria: 1.) Exons at least 1,500 bp in length (with one exception: *OMG*); 2.) Rapidly evolving with the raw percent divergence greater than or equal to that of *RAG1* for both deep and shallow splits (calculated by comparing *Anolis carolinensis* with *Gallus gallus* and *Python molurus*). *RAG1* has shown utility for reconstructing phylogenies at both deep and shallow levels (Groth and Barrowclough 1999) and has become one of the most commonly used nuclear markers in vertebrates (Barker et al. 2004; Townsend et al. 2004; Gruber et al. 2007; Hugall et al. 2007; Fuchs et al. 2011; Portik et al. 2011; Pyron 2011; Gamble et al. 2012; Harrington and Near 2012; Gartner et al. 2013); 3.) Presence in all amniote groups (confirmed in mammals, birds, and squamates with a few exceptions); and 4.) Single copy, with paralogs only allowed for duplications that predate the common ancestor minimally of squamates, (though in most markers it predates the common ancestor of all amniotes).

We identified many of the RELEC loci by querying the Ensembl database (Aken et al. 2017) for exons in *Anolis carolinensis* and other genomes. However, annotation errors are common on Ensembl especially for long exons (such as premature or incorrect boundaries or missing annotations) so we individually considered each locus to assess correct annotation and other criteria. We gathered additional exons by querying and inspecting all long mammalian exons on the Orthomam database (Douzery et al. 2014) and cross comparing the results on Ensembl for *Anolis* and across the amniote tree using BLAST searches. For example, for the 43 RELEC exons that are 4000 bp and longer, 27 were annotated by Ensembl on the *Anolis* genome and were easily identified, and 16 more were identified following searches on Orthomam and were not originally found because they were missing annotations or had errors on Ensembl. Using these methods, we are confident that we have gathered all or nearly all single copy exons that show this length and evolutionary rate and are also present among amniotes.

To avoid issues of gene duplications, we limited the set of loci in the RELEC dataset to those that were present in all amniote groups, and only included genes with known paralogs if duplications occurred on very deep lineages (e.g., for all mammals) and had been maintained. We also excluded some exons that present high levels of repeats that could bias phylogenetic analyses (e.g., *TTN*). Unlike other rapidly evolving regions of the genome that may mutate beyond recognition over deep time scales, exons are much more readily maintained across distantly related organisms, likely due to their direct functional role. For example, the major exons for the rapidly evolving genes *BRCA1* and *EXPH5* can be aligned and compared to examine evolution across all amniotes, whereas the intervening introns have accumulated substantial indels (see www.ensembl.org gene tree browser). After assembling the set of loci, we checked for presence and proper annotation of RELEC loci across 6 major tetrapod lineages using the chicken, saltwater crocodile, painted turtle, *Anolis*, human, and *Xenopus* genomes. We did this first by querying Ensembl gene names to extract annotation information for the longest exon. However, many genes did not have proper names on Ensembl, and for these we manually used tBLASTn searches on Ensembl to confirm presence. In these cases, we noted poor annotations when the exon or gene was either not annotated at all or had incorrect intron-exon boundaries, and determined the length of unannotated exons by searching for open reading frames surrounding the highest tBLASTn hit (Table S4). The final RELEC dataset includes 179 exons, representing 173 unique genes (6 genes with two exons each).

### Squamate Genomes

We queried 15 published and 2 unpublished squamate genomes to build the RELEC, UCE, and AHE datasets for comparison in this study. Assembled genomes were downloaded from NCBI(_*1*_), GigaScience (_*2*_), or unpublished (_*3*_). This includes the snakes *Python molurus_1_* (Castoe et al. 2013), *Boa constrictor_2_* (Bradnam et al. 2013), *Vipera berus_1_*, *Crotalus horridus_1_, C. mitchellii_1_* (Gilbert et al. 2014), *Deinagkistrodon actutus_2_* (Yin et al. 2016), *Protobothrops mucrosquamatus_1_* (Aird et al. 2017), *Ophiophagus hannah_1_* (Vonk et al. 2013), *Thamnophis sirtalis_1_* (Perry et al. 2018), and *Pantherophis guttatus_1_* (Ullate-Agote et al. 2014), and the lizards *Gekko japonicus_1_* (Liu et al. 2015), *Eublepharis macularius_2_* (Xiong et al. 2016), *Correlophus ciliatus_3_*, *Christinus marmoratus_3_*, *Dopasia gracilis_2_* (Song et al. 2015), *Pogona vitticeps_2_* (Georges et al. 2015), and *Anolis carolinensis_1_* (Alföldi et al. 2011). At the time of data acquisition, three more squamate species genomes were available for *Thamnophis elegans* (Vicoso et al. 2013)*, Sistrurus miliarus* (Vicoso et al. 2013), and *Sceloporus occidentalis* (Genomic Resources Development Consortium et al. 2015), though due to short read lengths and/or missing data, we chose not to use them (the purpose of these studies was not to acquire the high coverage or long read lengths needed for our purposes). The genomes of other squamates were published after our analyses were completed (*Shinisaurus crocodilurus*, Gao et al. 2017; *Salvator merianae*, Roscito et al. 2018; *Lacerta viridis* and *L. bilineata*, Kolora et al. 2019; *Varanus komodoensis*, Lind et al. 2019; *Zootoca vivipara*, Yurchenko et al. 2019) and we were not able to include them. The genomes we used encompass several divergent squamate lineages, but are primarily focused on snakes and geckos. Since geckos represent the sister clade to remaining squamates (Streicher and Wiens 2017), the dataset provides satisfactory depth for comparisons of the different methods.

### Data Generation and Alignments

We extracted and aligned sequences from the 17 genomes using customized scripts and using similar (though slightly adjusted) methods for each dataset. To extract both the RELEC and AHE loci from the genomes, we employed the BLAST+ command line tools (Camacho et al. 2009) to build local blast databases for each genome assembly. We developed the initial dataset for the RELEC loci with the genome of the gecko *Correlophus ciliatus*. After confirming the translation frame visually, we translated the sequences using the program Geneious v9.0.4 (Kearse et al. 2012), and then used the *tblastn* command from the BLAST+ toolkit to search for similar amino acid sequences in any frame in each genome. This method allowed for proper identification and extraction of orthologous exon regions even for rapidly evolving genes and the most divergent lineages. For the AHE dataset, we searched the genomes using the 389 version 2 vertebrate loci from Ruane et al. (2015). Since AHE markers include a combination of coding and noncoding regions, and because there is no detailed list of which markers are associated with specific genomic regions, we used the non-translated *blastn* search algorithm for each AHE search sequence.

BLAST searches for highly divergent sequences often produce multiple segmented hits for a single locus, rather than the entire length of the marker. In order to extract the entire orthologous sequence for each locus, we designed a script, Ortholog Assembly and Concatenation (OAC), in R v3.1.13 (R Core Team 2016) to combine coordinates of sequences that had more than one BLAST hit within close proximity on a single assembled contig, and extract and align the new sequence with a flanking region that was later trimmed to the initial query sequence (available at https://github.com/benrkarin/RELEC). This ensured that the maximum hit sequence length would be captured despite potential indels causing segmentation of BLAST hits to the same contig. Though on occasion some sequences were clearly split into multiple assembled contigs, we refrained from extracting the whole sequence to eliminate the possibility of unintentionally capturing paralogous sequences from different genomic regions. We indexed the genome assemblies using SAMtools (Li et al. 2009), and extracted the sequences based on the combined BLAST search coordinates for each marker with the *faidx* command. We then aligned individually extracted sequences by locus using MAFFT v7.130b (Katoh and Standley 2013), allowing the program to automatically determine sequence direction to accommodate for reversed sequences. All sequence alignments for RELEC and AHE were visually inspected in Geneious to confirm successful sequence capture and manually trimmed to reduce ragged ends. We conducted additional blast searches for any missing sequences, and to confirm we had captured the maximal sequence length available in the genomes. We conducted further blast searches of RELEC loci extracted from the *Correlophus ciliatus* genome against a combined transcriptome for *C. ciliatus* generated from six tissue types (eye, brain, tail, testis, ovary, whole embryo). We found transcripts corresponding to all RELEC loci in the transcriptome, confirming that RELEC genes are actually transcribed into mRNA.

For RELEC, we allowed incomplete alignments with less than the total 17 taxa for four exons: PKDREJ_B, which is absent in gekkotans and *Dopasia gracilis*, but is a very rapidly evolving and informative marker that will be useful in shallow-scale phylogenetic studies; TRANK1 exons 1 and 2, which are absent in the genomes of *Ophiophagus hannah*, *Thamnophis sirtalis*, and *Pantherophis guttatus*, and partially deleted in *Vipera berus*; and NAIP, which is absent from the *Pogona vitticeps* genome. For AHE, we only used completely sampled 17-taxa alignments.

For the UCE dataset, we followed the *phyluce* software package manual (Faircloth 2015) using the provided *Anolis* 5k probeset (available from https://github.com/faircloth-lab/phyluce), and extracted a flanking region of 300 bp for each UCE marker. This flanking region is comparable to mean aligned sequence lengths generated in empirical studies (e.g., Crawford et al. 2015: 820 bp; Grismer et al. 2016: 645 bp). For the UCE dataset, we used a complete matrix containing data for all 17 taxa for the main analysis (1,517 loci), but also generated datasets for loci with at least 15 taxa (additional 1,151 loci) and at least 9 taxa (additional 741 loci) for comparison (see Fig. S8, other results are comparable to the complete set and are not shown).

We used reciprocal BLAST searches to assess overlap between datasets. We found that 17 AHE loci match UCE loci; none of the RELEC loci match any UCE loci; and 28 RELEC loci match AHE loci (4 of which are from Wiens et al. 2012). Though for these overlapping regions it is important to note that RELEC targets the entire exon rather than a portion of it or the flanking intron.

### Phylogenetic Analyses

We conducted summary-coalescent species tree analyses using ASTRAL-III (Zhang et al. 2018) based on sets of 100 maximum likelihood bootstrap replicates generated on each locus individually in RAxML v8.1.15 (Stamatakis 2014), and specified all four of the gekkotans as the rooting taxa. We chose the GTRGAMMA model for all analyses, as Stamatakis (2015) suggests that other models may be inappropriate for datasets with relatively few taxa such as this one. We also generated gene trees for RELEC loci translated to amino acid sequences in RAxML, allowing the program to choose the model with the PROTGAMMAAUTO setting. We conducted a final ASTRAL analysis on a combined dataset of all AHE, UCE, and RELEC loci (2,016 loci after excluding any overlapping loci). We also conducted parallel analyses in IQTree (Nguyen et al. 2015) allowing for automatic model selection (Kalyaanamoorthy et al. 2017) and recovered nearly identical results for gene tree and ASTRAL estimates (not shown).

Concatenated datasets and partition files were generated using the perl program, BeforePhylo v0.9.0 (Zhu 2014), and concatenated maximum likelihood trees were generated with 1000 bootstrap replicates with RAxML, under the same rooting and model settings as the gene trees. To maintain consistency across datasets, we partitioned the concatenated analyses by each locus, without specifying different partitions for codon positions in the RELEC dataset. To improve computing time, we ran the concatenated analyses on the CIPRES Science Gateway (Miller et al. 2010).

We analyzed concordance and accuracy of the maximum likelihood gene trees in R by incorporating functions from the *ape* (Paradis et al. 2004) and *phangorn* (Schliep 2011) packages to compute the proportion of the highest scoring maximum likelihood trees in each dataset that display a particular node in the species tree. This is similar to the gene concordance factor that can be estimated in IQTree (Minh et al. 2018) (results in Table S3), but our technique uses a different rooting method that allows for a proportional value to be calculated at every node, whereas the concordance factor excludes some nodes. We also examined the Robinson-Foulds (RF) distance between the best scoring individual gene trees and the species tree. RF distances quantify the accuracy of gene trees, with values of 0 corresponding to an exact match, a value of 2 corresponding to one node difference, and so on. To examine the accuracy of gene trees at a finer scale, we conducted the same analysis restricted to the eight colubroid snakes in the tree. We assessed the robustness of gene trees to correctly build the species tree (using the dataset-specific ASTRAL species tree to reduce bias) by iteratively running ASTRAL on randomly selected jackknife replicate bins of varying numbers of gene trees from two gene trees to the full number in each dataset using a python script provided by Daniel Portik.

We used the multispecies coalescent to estimate species trees from the complete set of 179 RELEC loci using StarBEAST2 (Ogilvie et al. 2017), part of the BEAST2 package (Bouckaert et al. 2014). This is a new implementation of the *BEAST method (Heled and Drummond 2010) but is considerably faster and enables the use of dozens or hundreds of loci. We used the analytical population size integration, with a strict molecular clocks and general time reversible (GTR) plus gamma model for each locus. We initiated two analyzes for 50 million generations each with a pre-burnin of 1 million generations, sampling every 5000 generations. The posterior distribution of post-burnin species trees was visualized using DensiTree v2.2.2 (Bouckaert and Heled 2014).

### Summary Statistics

We computed general summary statistics (see Table S1) for each marker in R using the packages *ape* (Paradis et al. 2004), *phyloch* (Heibl 2016), and *adephylo* (Jombart and Dray 2016). These included the GC content across all sites, GC content at codon position 3 (GC3), average pairwise identity for alignments, proportion of variable and parsimony informative sites, and raw pairwise genetic distance between *Gekko japonicus* and *Anolis carolinensis* as a rough relative measure of evolutionary rate (Table S1). We also calculated the number of segregating sites (Fig. S5), and assessed the size and width of alignment gaps (Fig. S6).

### Phylogenetic Informativeness and Substitution Saturation

We profiled the phylogenetic informativeness (Townsend 2007) of each marker using the web based tool PhyDesign (López-Giráldez and Townsend 2011), specifying the DNArates algorithm. This method provides an estimate of the relative power of each locus to resolve a given node in the tree. The program requires a timetree as input, so we transformed the output ASTRAL phylogeny (AHE, UCE, and RELEC combined dataset topology) into a timetree using the *ape* package and *ScalePhylo* (Hunt 2011) script in R. *ScalePhylo* allowed us to force a time-calibration on the nodes of the tree based on input divergence dates, which we acquired from the date estimate for each pair of taxa from http://timetree.org/ (Hedges et al. 2015). We downloaded the output from the PhyDesign web portal, and plotted the measures of phylogenetic informativeness for each dataset separately in R to allow for comparison between datasets. We visually examined substitution saturation in the RELEC loci by generating plots of raw sequence divergence at divergence date estimates from the http://timetree.org/ database. We used the output in R to compare how maximum informativeness scales with alignment length, and also the time of maximum informativeness. We also scaled the curves by alignment length in order to assess the informativeness independent of length (results shown in Fig. S7)

### Biological Processes

We sought to investigate if RELEC loci are subject to unique biological processes that contribute to their rapid evolution and maintenance of long exon length. We compared the expression levels across human tissues using the transcript abundance categorization of Fagerberg et al. (2014), and also compared expression of RELEC genes relative to the background within individual tissue types. We searched the Swiss-Prot database (http://www.uniprot.org) for the number of GO terms associated with each protein for *Homo sapiens* from any of the three GO categories (biological process, molecular function and cellular component). This gives an approximation of the breadth of functional utility for each protein, and may potentially influence the rate of molecular evolution. We also searched the BioGRID database (http://thebiogrid.org) for the number of unique protein interactors associated with each protein in *Homo sapiens*, for which the data is most extensive. Finally, we looked for GO term enrichment using the web server for the PANTHER 14.1 classification system (http://pantherdb.org).

## Data Availability

Scripts, sequence alignments, and other resources for mammals, birds, and squamates are available at https://github.com/benrkarin/RELEC.

## Supplementary Material

Supplementary data, including figures, tables, and alignments are available at Molecular Biology and Evolution online. Fig. S1: DensiTree plot of the post-burnin species trees from the StarBEAST2 analyses. Fig. S2: Concatenated maximum likelihood trees. Fig. S3: RELEC ASTRAL and concatenated translated amino acid trees. Fig. S4: Saturation plots of all loci with divergence time (Ma) on the x-axis and raw pairwise distance is the y-axis. Fig. S5: Proportion of segregating sites in alignments of each dataset. Fig. S6. Proportion of alignments as gaps, mean width of gaps, and the relationship between proportion of gaps and the proportion of parsimony informative sites. Fig. S7. Phylogenetic informativeness scaled by alignment length. Fig. S8. Comparison of increasing numbers of UCE loci of differing completeness (9-14 taxa, 15-16 taxa, all 17 taxa). Table S1: Description of RELEC loci, including summary statistics, number of GO processes, and number of Interaction terms. Table S2: Genomic position and gene name of *Anolis carolinensis* AHE version 2 vertebrate loci (probes from Ruane et al. 2015). Table S3. Gene and site concordance factors calculated in IQTree. Table S4. Ensembl stable IDs, exon position, and exon length for RELEC loci in tetrapod lineages (source data for Fig. 1).

## Acknowledgements

We are grateful for permission to use the photographs in Fig. 1 from Sean Harrington (*C. mitchellii*), Alex Krohn (*T. sirtalis*, Patrick Campbell (*B. constrictor*), Hans Breuer (*P. mucrosquamatus*), 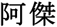 (*D. acutus*), Takuya Morihisa (*G. japonicas*), Sandesh Kadur/Felis Images (*D. gracilis*), Warren Photographic (*P. molurus*, *P. guttatus*, *V. berus*), Tom Charlton (*O. hannah*), Todd Pierson from calphotos.berkeley.edu CC BY-NC-SA 3.0 (*A. carolinensis*), TG (*C. ciliatus*, *E. macularius*, *C. marmoratus*, *P. vitticeps*) and BRK (*C. horridus*). We thank Nick Crawford for input on developing scripts. We thank Dr. Jimmy McGuire, Dr. Rauri Bowie, and their lab groups for providing critical input on the manuscript. We thank Daniel Portik for use of a python script to run the gene tree jackknife analysis. We especially want to acknowledge Margarita Metallinou, who provided valuable insight into this project in its initial stages. The U.S. National Science Foundation funded gecko genome sequencing (NSF IOS1146820) and partial support for TG (NSF DEB1657662).

