## Supplementary figures and images for "Optimizing Phylogenomics with Rapidly Evolving Long Exons: Comparison with Anchored Hybrid Enrichment and Ultraconserved Elements"

### Fig. S4

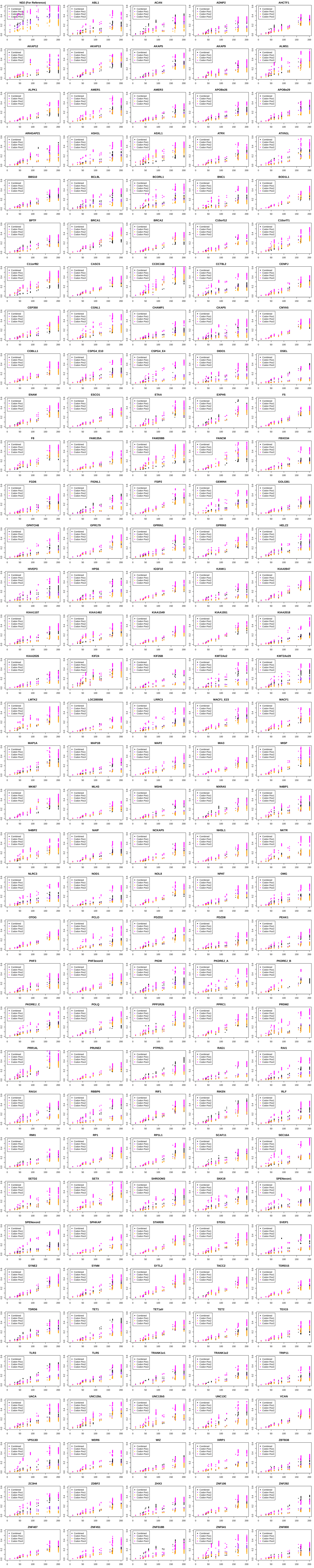
