## Supplemental Data 1 for "Optimizing Phylogenomics with Rapidly Evolving Long Exons: Comparison with Anchored Hybrid Enrichment and Ultraconserved Elements"

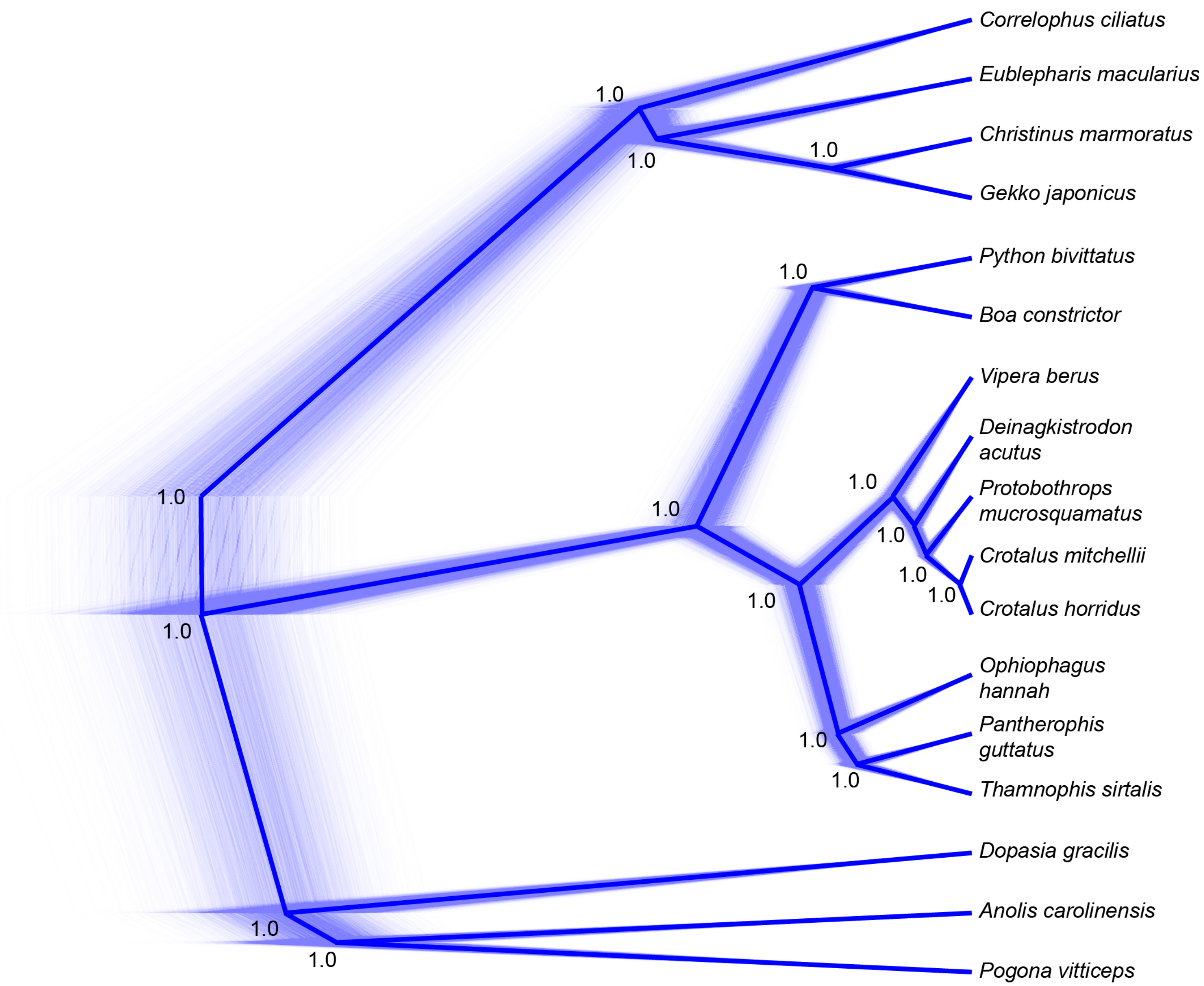


**Fig. S1.** DensiTree plot of the post-burnin species trees from the StarBEAST2 analyses. Posterior probabilities are shown at each node.

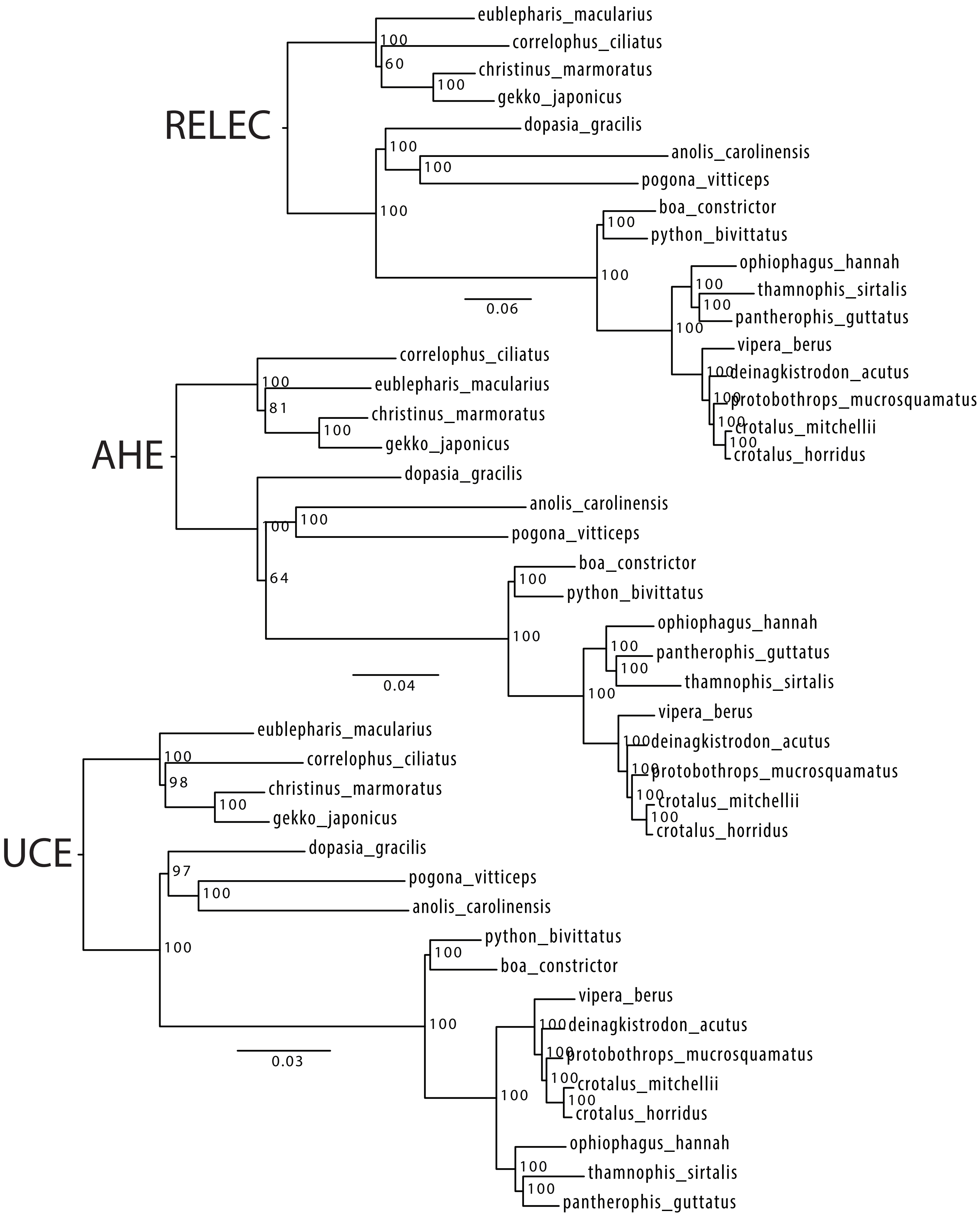


**Fig. S2.** Concatenated and ASTRAL species tree of RELEC translated amino acid dataset.

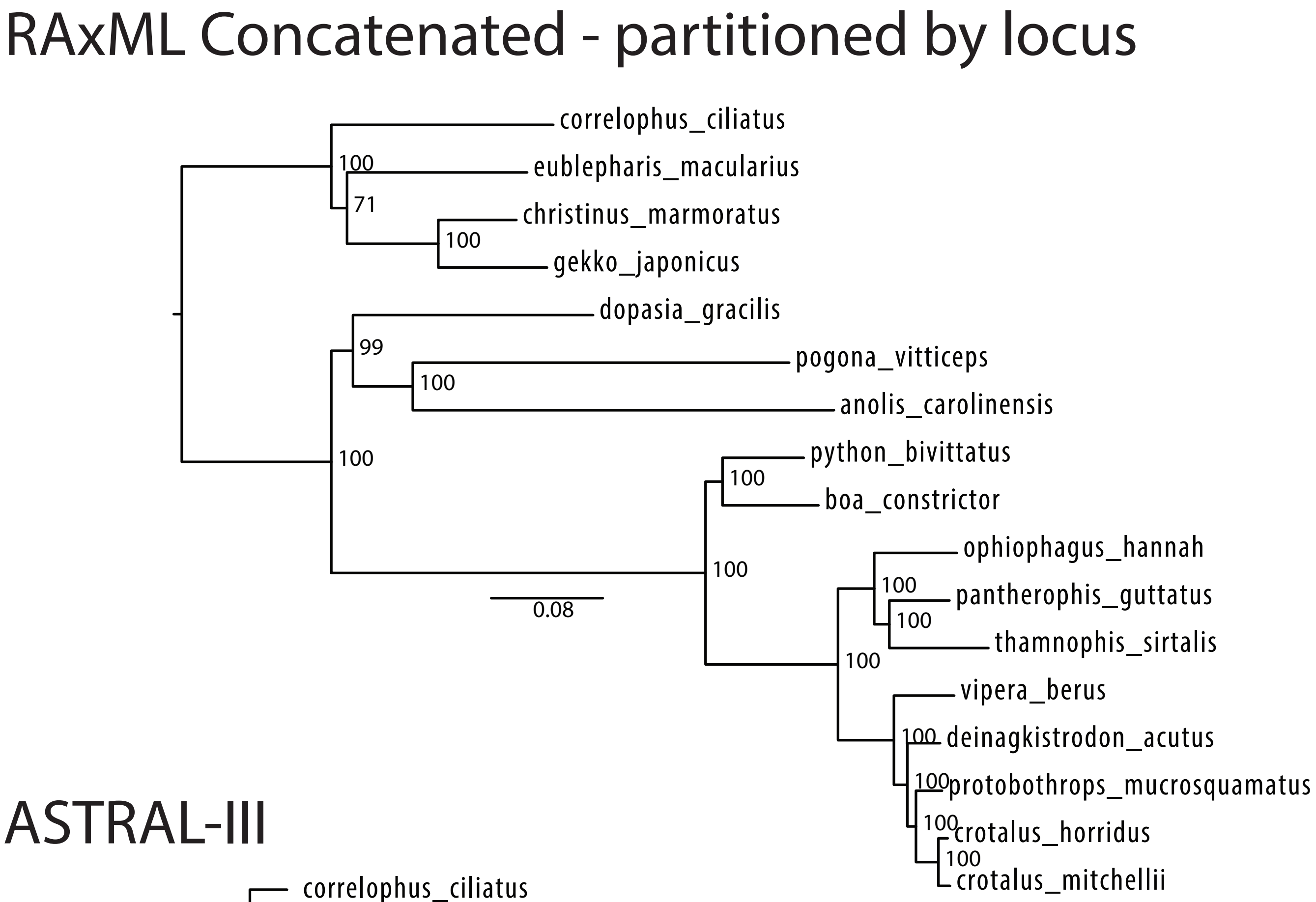


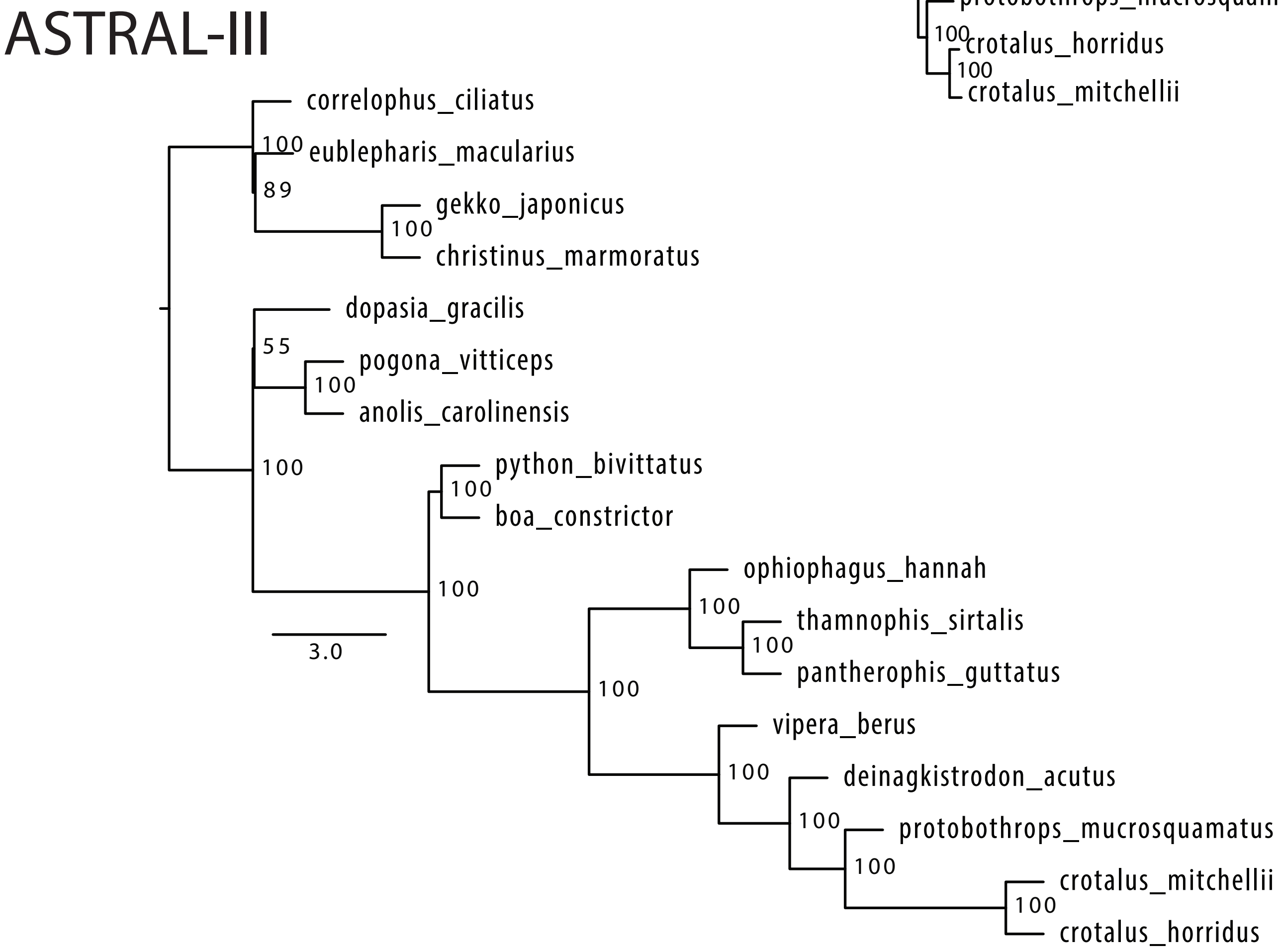


**Fig. S3.** Concatenated and ASTRAL species tree of RELEC translated amino acid dataset.

**Separate File**

**Fig. S4.** Saturation Plots of all loci. Divergence time (Ma) is the x-axis, and raw pairwise distance is the y-axis.

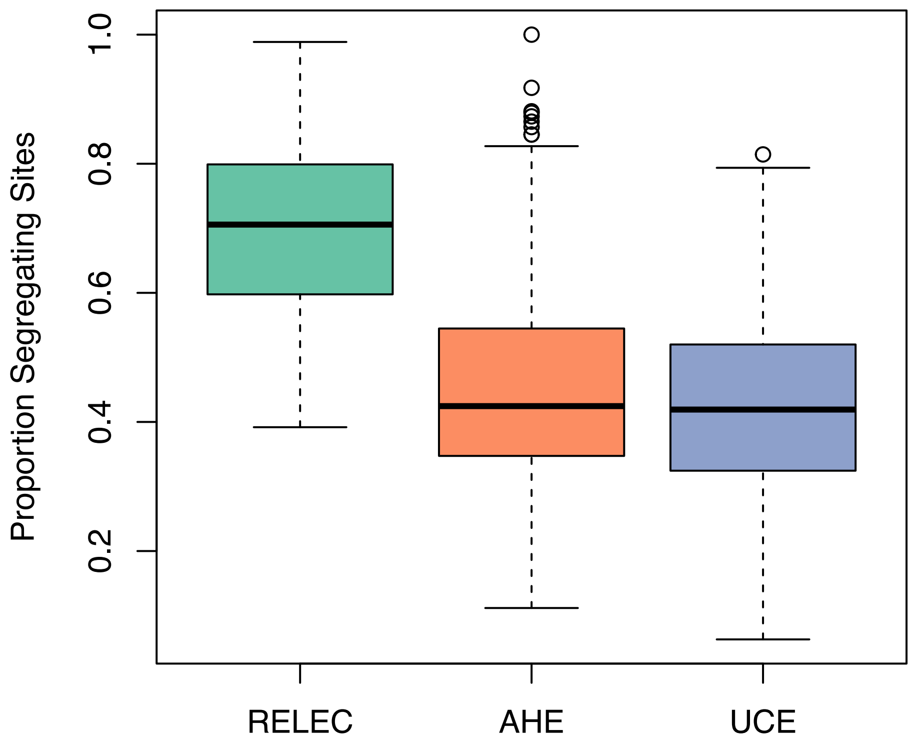


**Fig. S5.** Proportion of segregating sites within RELEC, AHE, and UCE alignments.

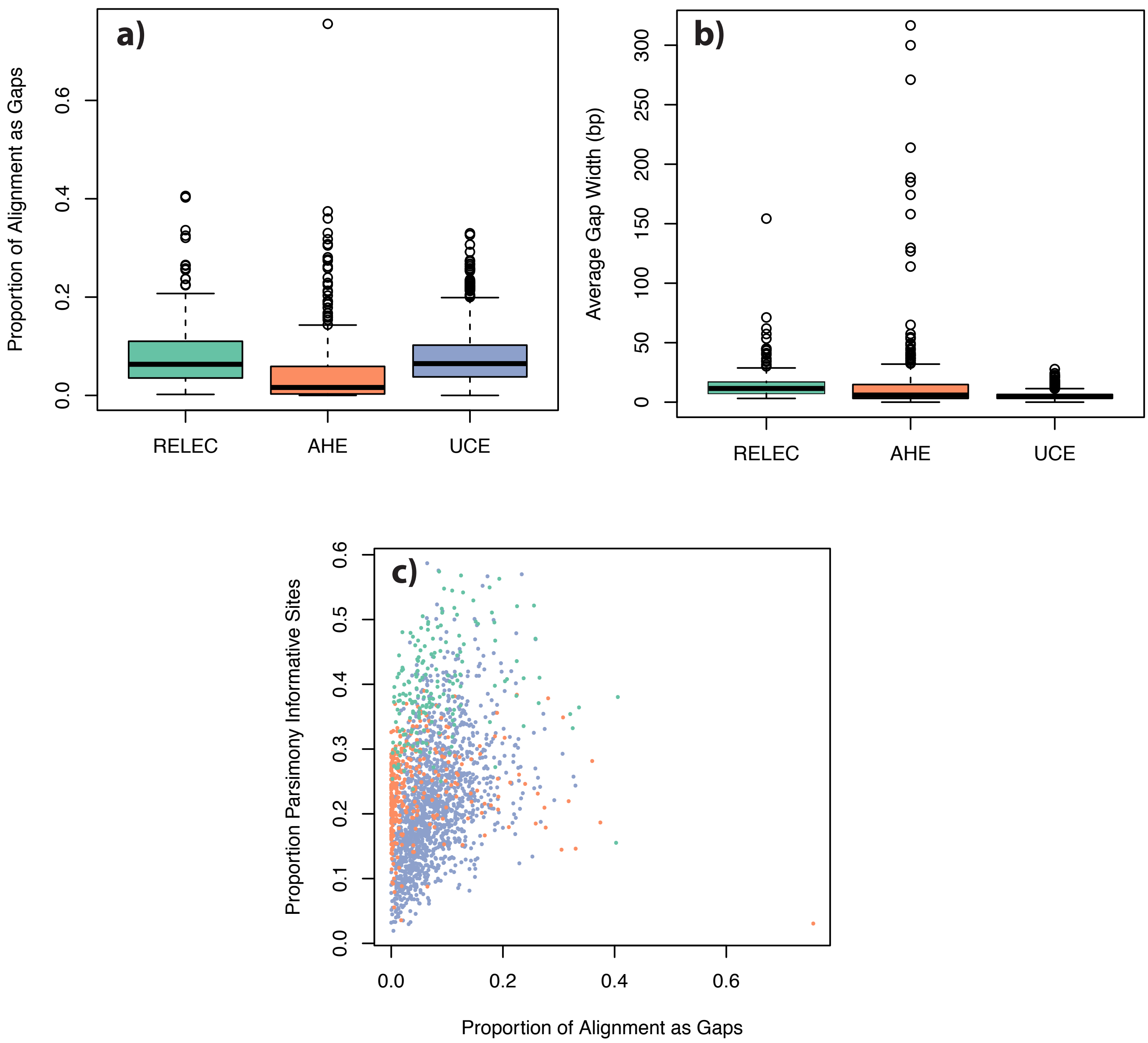


**Fig. S6.** Comparison of RELEC, AHE and UCE alignments for: a) the proportion of the alignent as gaps; b) the average gap width within locus alignments; and c) the relationship between the proportion of gaps and the proportion of parsimony informative sites in each alignment. RELEC loci show more PIS relative to the proportion of gaps, suggesting that they may provide more information content with less alignment issues. Color coding of (c) matches boxplots.

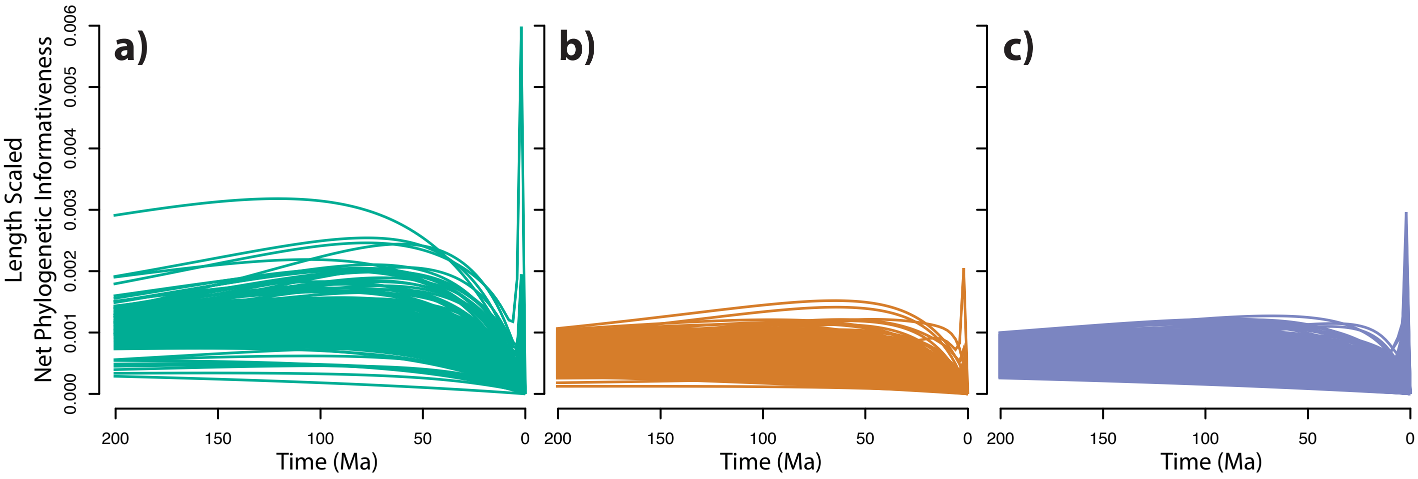


**Fig. S7.** Phylogenetic informativeness for (a) RELEC, (b) AHE, and (c) UCE loci, scaled by alignment length.

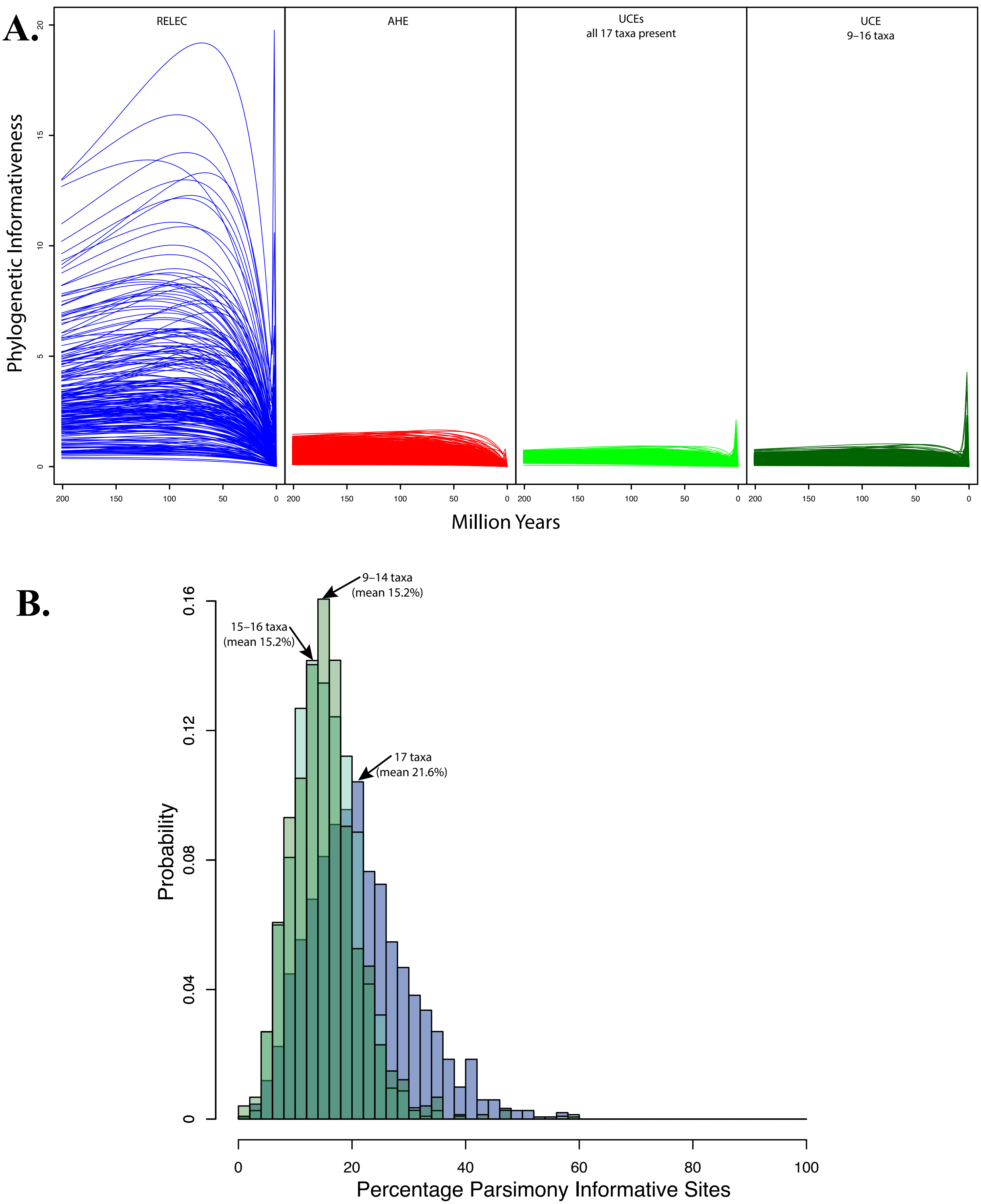


**Fig. S8.** Comparison of phylognetic informativeness (A) and the percentage of parismony informative sites (B) in UCE datasets that are complete with 17 taxa to those with differing amounts of missing data.
